# Endogenous gene editing of alveolar organoids reveals that expression of pathogenic variant SFTPC-I73T disrupts endosomal function, epithelial polarity and wound healing

**DOI:** 10.1101/2025.07.22.665497

**Authors:** Eimear N. Rutherford, Dawei Sun, Kyungtae Lim, James R. Edgar, Lydia E. Matesic, Stefan J. Marciniak, Emma L. Rawlins, Jennifer A. Dickens

**Affiliations:** Cambridge Institute for Medical Research, Cambridge, CB2 0XY, UK; Gurdon Institute, University of Cambridge, Cambridge CB2 1QN, UK; Department of Physiology, Development and Neuroscience, University of Cambridge, Cambridge CB2 3DY, UK; Broad Institute of Massachusetts Institute of Technology and Harvard, Cambridge, MA 02142, USA; Department of Life Sciences, Korea University, 145 Anam-Ro, Seoungbuk-Gu, Seoul 02841, South Korea; Department of Pathology, University of Cambridge, Cambridge, CB2 1QP, UK; Department of Biological Sciences, University of South Carolina, 715 Sumter St., Columbia, SC 29208, USA; Royal Papworth Hospital, Papworth Road, Trumpington, CB2 0AY

## Abstract

**Background:** Idiopathic pulmonary fibrosis is a fatal lung disease of progressive lung parenchymal scarring caused by the aberrant response of an alveolar epithelium repeatedly exposed to injury. Understanding epithelial dysfunction has been hampered by the lack of physiological alveolar type 2 (AT2) cell models and defined disease triggers. Monogenic forms of familial pulmonary fibrosis (FPF) caused by toxic gain-of-function variants provide an opportunity to investigate early pathogenic events. One such variant, surfactant protein C (SFTPC)-I73T, abnormally localises within AT2 cells and causes their dysfunction.

**Methods:** We used base editing of fetal lung-derived AT2 (fdAT2) organoids to create a heterozygous disease model of endogenous SFTPC-I73T expression. We also created an inducible overexpression system to interrogate temporal changes associated with SFTPC-I73T expression. We cultured fdAT2 both in 3D culture and at air-liquid interface to understand the importance of polarity cues and air exposure on disease phenotypes.

**Results:** In our heterozygous endogenous expression system, we found that fdAT2 expressing SFTPC-I73T grew without a lumen and were unable to correctly polarise. SFTPC-I73T accumulated with time and caused gross enlargement of early endosomes, preventing correct apico-basal trafficking of multiple endosomally trafficked cargoes including polarity markers and cell adhesion proteins. This phenotype was exacerbated by air exposure and led to loss of epithelial monolayer integrity and abnormal wound healing after injury.

**Conclusion:** Using endogenous gene editing for the first time in differentiated alveolar organoids, we have demonstrated that the pathogenic effects of SFTPC-I73T are mediated through endosomal dysfunction and abnormal epithelial organisation. This has important implications for AT2 function *in vivo*.

## Introduction

Idiopathic pulmonary fibrosis (IPF) is a devastating disease of relentlessly progressive lung parenchymal scarring which leads to impaired gas exchange and premature death^1^. Current therapies, which target downstream pathogenic pathways, are incompletely effective and poorly tolerated^2,3^. Thus, there is an urgent unmet need to understand and target early pathogenic events.

The alveolar epithelium is now understood to be key in disease initiation; this is in part evidenced by the observation that pathogenic variants of AT2-specific surfactant-related genes are responsible for some monogenic inherited forms of pulmonary fibrosis^4^. Repeated injury to the alveolar epithelium, often from inhaled toxins, provokes a change in the behaviour of vulnerable alveolar type 2 (AT2) cells. In health, they act as alveolar-resident stem cells, differentiating to alveolar type 1 (AT1) cells and migrating to repair injured epithelium^5^. In IPF, their phenotype is fundamentally altered. Their ability to differentiate, proliferate and migrate are impaired and, along with an altered secretome, their phenotype switches from pro-repair to pro-fibrotic. This triggers downstream pathogenic events which lead to pathological lung parenchymal remodelling and fibrosis^6^. Understanding how AT2 dysfunction triggers disease is crucial for therapeutic targeting but has been challenging both due to the absence of a single exogenous disease trigger and a lack of phenotypically stable *in vitro* AT2 models.

Familial pulmonary fibrosis (FPF) caused by single gene variants provide a unique opportunity to study triggers of AT2 dysfunction. The variants in surfactant-biology associated genes which cause intrinsic AT2 dysfunction through toxic gain-of-function mechanisms are particularly valuable. Autosomal dominant mutations in surfactant protein C (SFTPC) cause FPF^7^. Disease can present from neonatehood through to adulthood, suggesting additional genetic, epigenetic or exogenous modifiers are important^8^. Many variants misfold, leading to ER retention and ER stress, something seen in alveolar epithelium in both FPF and sporadic disease^9,10^. The commonest pathogenic variant, SFTPC-I73T, rather mislocalises to the plasma membrane^11–16^. We and others have shown that is because SFTPC-I73T cannot access late trafficking compartments due to failure of ubiquitination and instead is recycled back to the plasma membrane^15,17,18^.

The mechanisms by which this protein accumulation leads to AT2 dysfunction are still incompletely understood, though it has been observed in both cell and animal models that AT2 cells expressing SFTPC-I73T show impaired autophagic flux and mitophagy. Transcriptomic and mouse data also support SFTPC-I73T expression creating a pro-inflammatory phenotype^14,16^. Elucidating the pathways driving these phenotypes is critical, as they may converge with mechanisms implicated in sporadic pulmonary fibrosis, such as those triggered by ER stress-inducing SFTPC variants, and identify shared targets for therapeutic intervention.

Recent advances in human *in vitro* AT2 culture have enabled more refined modelling of AT2-specific disease mechanisms. Expandable, phenotypically stable AT2-like cells can be derived from induced pluripotent stem cells (iAT2s)^19^ or lung epithelial tip progenitor cells from fetal lung tissue (fdAT2s)^17,20^. Both systems express key AT2 markers and are capable of producing and processing pulmonary surfactant, making them valuable tools for probing cell-intrinsic dysfunction. While fdAT2 lines have been manipulated using integrated CRISPR interference (CRISPRi) to modulate gene expression at the transcriptional level^17^, endogenous gene editing or direct modelling of disease-causing mutations in these cells has not yet been achieved. iAT2s can be genetically manipulated as iPSCs before differentiation to generate isogenic lines; however, this approach is both technically and temporally arduous and typically relies on access to patient samples from those with very rare conditions^16^.

To overcome these methodological constraints and tackle unanswered mechanistic questions, we report here for the first time endogenous base editing of differentiated fdAT2 organoids enabling rapid generation of a heterozygous SFTPC-I73T model. We characterise these cells both when grown as organoids and then at a more physiological air-liquid interface (ALI). We find that SFTPC-I73T accumulation causes early endosomal engorgement leading to loss of polarity and abnormal localisation of other endosomally trafficked cargoes, including cell adhesion proteins. This culminates in loss of epithelial integrity and impaired repair following injury.

By developing a novel way to model disease in alveolar organoids, this work defines a previously unrecognised mechanism by which SFTPC-I73T disrupts AT2 homeostasis and establishes a tractable platform to study intrinsic epithelial dysfunction in pulmonary fibrosis. Our findings underscore the importance of endosomal trafficking in maintaining AT2 cell function and implicate its disruption as an initiating event in fibrotic lung disease.

## Results

To interrogate the mechanism of SFTPC-I73T induced alveolar epithelial cell dysfunction, we generated a T>C substitution at c.218 of the human *SFTPC* gene in fdAT2. As there is no suitably located classical “NGG” PAM sequence to facilitate this edit, we employed a near PAM-less SpRY-Cas9 along with an adenine base editor (ABE)^21^ to induce an A>G substitution on the complementary strand (Fig. 1A-C). gRNAs were designed by selecting sequences adjacent to NGN or NAN PAM motifs near the target site and ensuring that the target adenine was positioned within the optimal editing window of the ABE (4th to 8th nucleotides of the protospacer sequence)^21^ (Fig. 1B).

**Fig 1.**
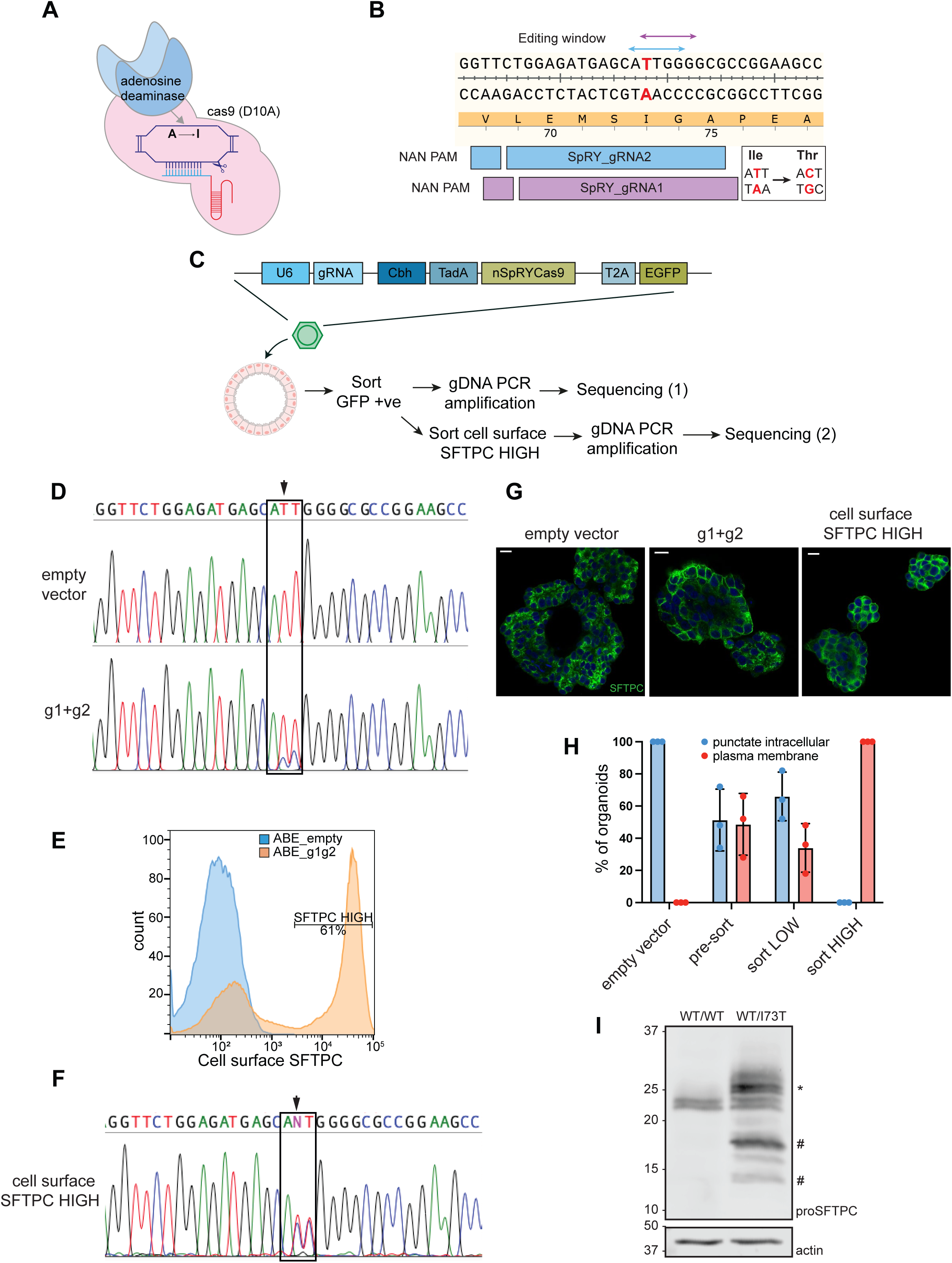
Generation of heterozygous endogenous SFTPC-I73T organoid model using an adenine base editor. (A) Schematic of adenine base editor containing a modified SpRY Cas9 (D10A) nickase and adenosine deaminase which converts targeted adenine to inosine, resulting in a permanent A-to-G conversion during DNA replication. (B) Location of gRNA used to introduce the A-to-G mutation on the complementary strand to the T-to-C base change required for the I73T mutation. The double ended arrows define the editing window for each guide. (C) Schematic of the expression plasmid and base editing strategy. (D) Sanger sequencing chromatogram showing target region of SFTPC gRNA after sorting for GFP positive cells (empty vector, base editor containing no guides; g1g2, base editor containing two guides targeting SFTPC). (E) Flow cytometry histogram of cell surface SFTPC in GFP positive cells as measured by C-terminal domain antibody. Cells with high cell surface SFTPC were sorted and expanded. (F) Sanger sequencing chromatogram showing target region of the SFTPC gRNA after sort for cells with increased SFTPC at cell surface. (G) Immunofluorescence imaging of proSFTPC in organoids transduced with an empty vector (left panel) or base editor (middle and right panels) pre- and post-sort. Scale bar 10µm. Quantified in (H), n=100 for each of n=3 biologically independent lines. (I) Immunoblotting for proSFTPC in edited and control fdAT2. *higher molecular weight full length SFTPC-I73T caused by O-glycosylation; #partially and fully C-terminally cleaved intermediates.

Organoids were transduced with lentiviruses containing 2 different sgRNAs targeting *SFTPC* to maximise the likelihood of successful editing (ABE_g1g2) (Fig. 1B). Control organoids were transduced with vectors lacking a gRNA sequence (“empty vector”). Three biologically independent organoid lines were generated. Transduced cells were sorted by GFP positivity (empty vector; ∼30% GFP +ve, SFTPC gRNA; <10% GFP +ve) and replated for recovery and expansion. Sanger sequencing revealed successful editing of c.218 in approximately 25% of *SFTPC* alleles (Fig. 1D, arrow). Off-target editing of the adjacent adenine (c.219) was observed but was functionally insignificant, producing a silent mutation (Fig. 1B&D). The edited population was isolated by sorting for fdAT2 enriched for cell surface-resident SFTPC, the hallmark of I73T expression (Fig. 1E)^12,14,15,22^. Subsequent sequencing indicated an almost exact 50% T>C edit (Fig. 1F and suppl Fig. 1A, arrows) and immunofluorescence microscopy demonstrated redistribution of SFTPC to the cell surface in 100% of organoids grown from these sorted populations (Fig. 1G&H). These data suggest successful generation of a pure population of SFTPC-WT/I73T heterozygous fdAT2 (hereafter “I73T-fdAT2”), suitable for modelling autosomal dominant SFTPC-I73T-associated disease. Immunoblotting of I73T-fdAT2 revealed both a band pattern that phenocopied the unedited organoids plus additional bands characteristic of the altered post-translational processing of SFTPC-I73T (*higher molecular weight species due to O-glycosylation; #accumulation of partially and fully C-terminally cleaved intermediates) (Fig. 1I and suppl Fig. 1B)^12,15^.

Importantly, genotypic stability was maintained across all lines to at least passage 10 (organoids were not used beyond this) and following cryopreservation-thawing, reinforcing our conclusion that the I73T-fdAT2 are a pure heterozygously edited population (suppl Fig. 2A). Despite this, phenotypic drift was observed at later passage in one I73T-fdAT2 line, but not in its control counterpart, in which cell surface SFTPC was dramatically depleted (suppl Fig. 2B, line 3) likely due to reduced SFTPC expression (suppl Fig. 2C). Although not further formally interrogated, we hypothesise that these changes may reflect a compensatory response to the toxic gain-of-function mutant. These observations highlight the importance of phenotypic characterisation throughout the culture period, as genotypic stability alone does not ensure model fidelity. Having identified this issue, all experiments were undertaken at passage 1-5.

We next examined the impact of heterozygous SFTPC-I73T expression on fdAT2 organoid appearance. As expected, control organoids grew with a folded morphology and expressed SFTPC which was largely localised to intracellular punctae. In contrast, the I73T-fdAT2 appeared macroscopically less well organised and aberrantly accumulated SFTPC at the plasma membrane, reproducing previous observations in PSC-iAT2, *in vivo* mouse models, and patient tissue (Fig. 2A&B)^14,16^. Strikingly, the loss of organisation in I73T-fdAT2 was demonstrated to reflect loss of lumen formation, with only a minority able to form a central lumen (Fig. 2C). This suggests a loss of apicobasal polarity, previously noted in PSC-iAT2^16^ and reminiscent of the morphological changes described in embryonic mouse lungs expressing HA-SFTPC-I73T^14^. In contrast to controls, I73T-fdAT2 displayed non-discriminate localisation of AT2 apical protein HTII-280, epithelial basolateral protein EpCAM, and adherens junction protein E-cadherin at the cell surface (Fig. 2D). Muc1 staining in I73T-fdAT2 suggested the presence of small, sometimes multiple non-central lumens (arrowheads) indicative of abnormal lumenogenesis. This has previously been associated with intracellular accumulation of apical proteins^23^. Perturbed lumenogenesis resulting in multiple-lumen and no-lumen phenotypes can arise from interference with the regulation of polarity complexes^24–28^. Staining of Crumbs protein homolog 3 (CRB3), which plays a crucial role in the establishment and maintenance of the apical membrane and epithelial cell polarity^29–34^ and which has apical punctate localisation in human adult AT2 cells^35^, was also disrupted with evidence of marked intracellular accumulation and redistribution to the cell surface in I73T-fdAT2 (Fig. 2E).

**Fig 2.**
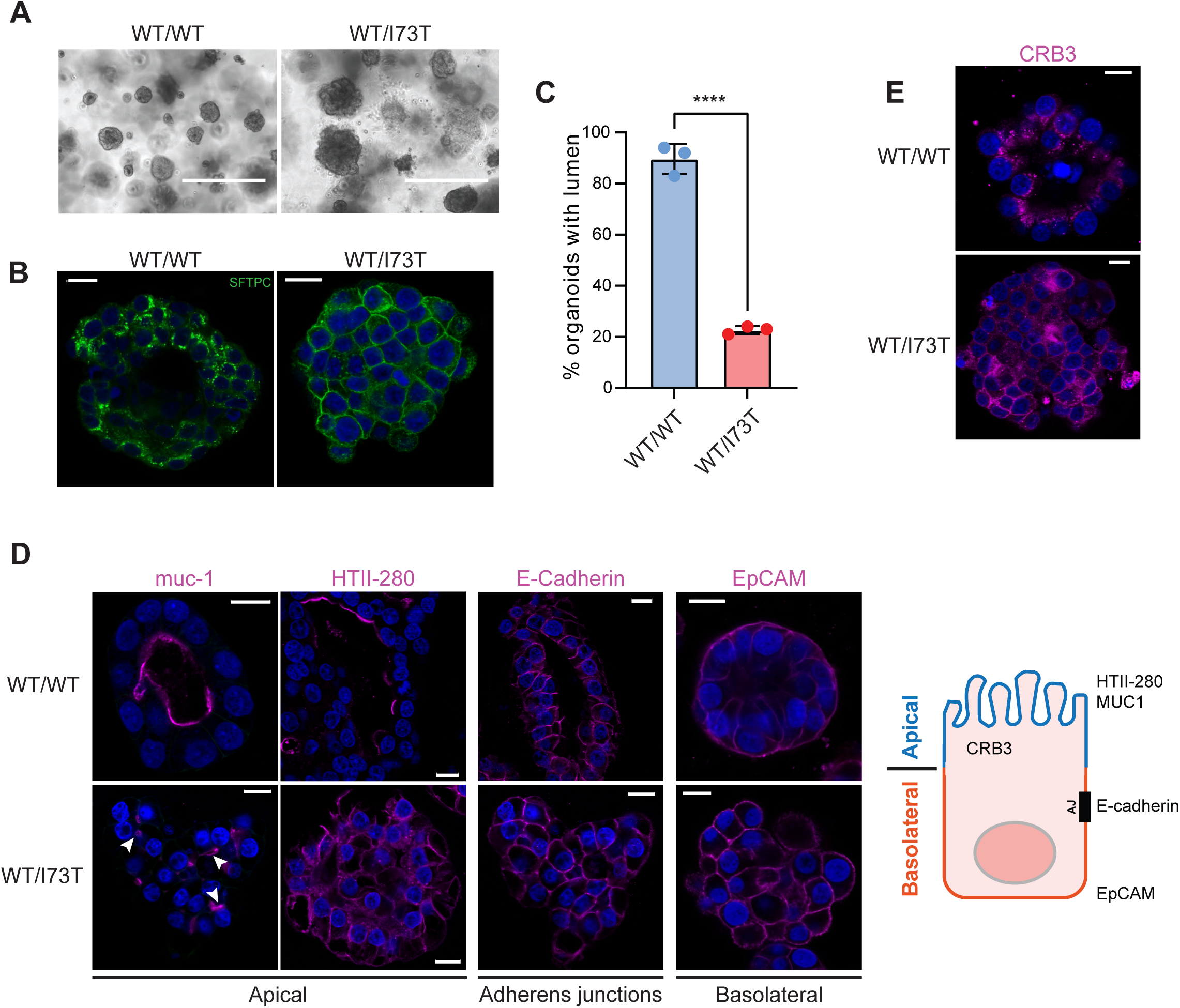
SFTPC-I73T expression disrupts lumenogenesis and epithelial polarity. (A) Brightfield and (B) proSFTPC immunofluorescence imaging of control and base-edited organoids. Scale bar, 400µm (brightfield images), 10µm (confocal images). (C) Percentage of organoids with lumen present in SFTPC-WT/WT and WT/I73T fdAT2. n=100 for each for each of n=3 biologically independent lines. Unpaired t-test. (D) Immunofluorescence imaging of muc-1, HTII-280, E-cadherin and EpCam with schematic of expected subcellular localisations. Scale bar, 10µm. (E) Immunofluorescence imaging of apical polarity marker CRB3. Scale bar, 10µm. ****=p<0.0001.

Establishment and maintenance of epithelial cell polarity relies on functional endosomal sorting of apical and basal cargoes, particularly cell polarity complexes and cell adhesion molecules^36,37^. We and others have shown that if SFTPC is unable to traffic into intraluminal vesicles of multivesicular bodies (MVBs), partially cleaved intermediates instead accumulate in early and recycling endosomes and at the plasma membrane^13,15^. To determine the effect of SFTPC-I73T expression on early endosomes in fdAT2, we stained organoids for the early endosome marker EEA1. As expected, SFTPC-WT was localised in small intracellular puncta which did not co-localise with EEA1, consistent with its trafficking to later compartments (Fig. 3A). In contrast, SFTPC was found in EEA1 positive compartments in I73T-fdAT2, which were significantly enlarged when compared with controls (Fig. 3A&B). Notably, early endosomes which did not contain SFTPC were also enlarged (arrowhead; Fig. 3A) and EEA1 was also partially redistributed to the cell surface, suggesting a functional effect on endosomal trafficking not limited only to SFTPC-containing vesicles.

**Fig 3.**
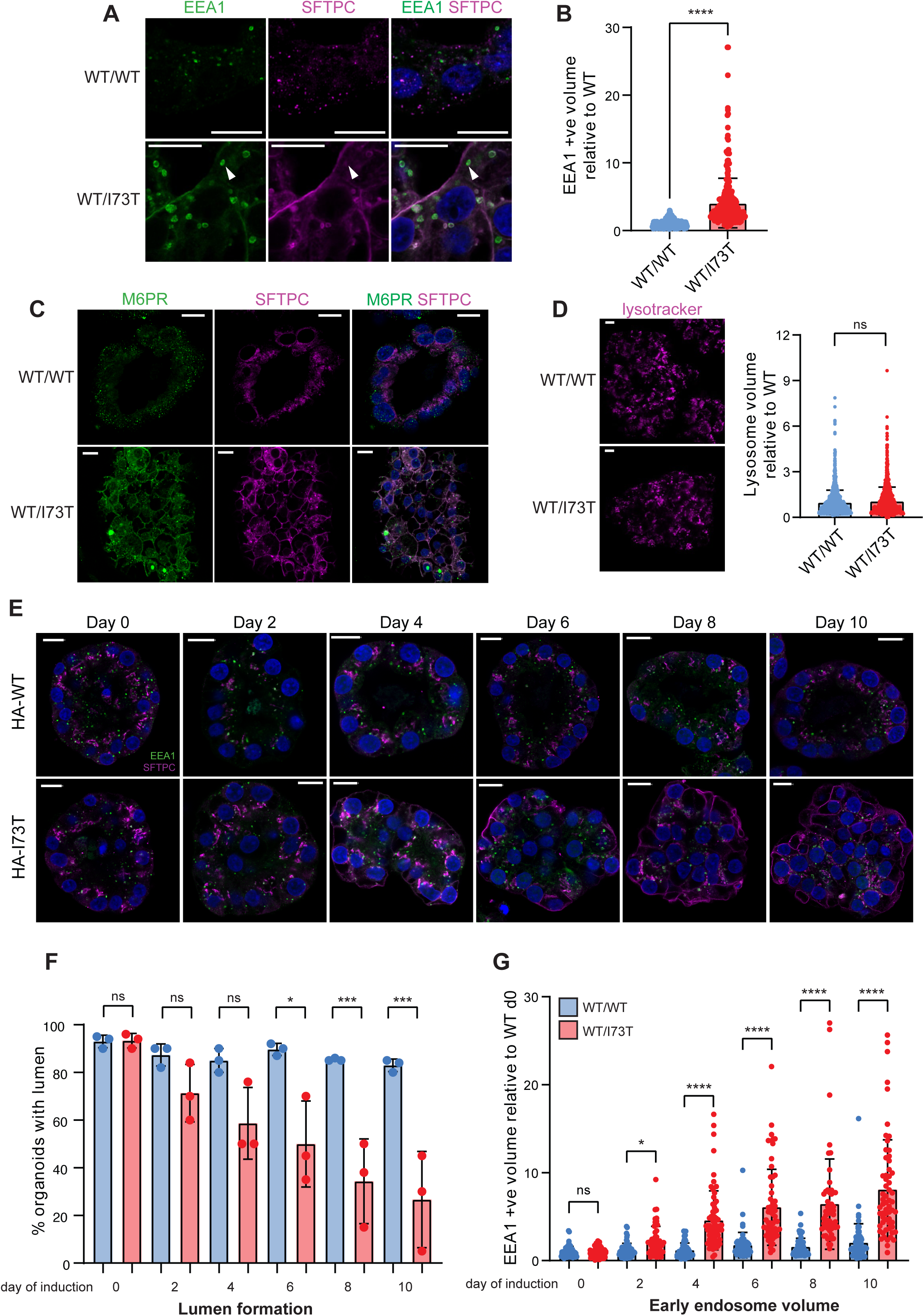
Endosomal enlargement due to SFTPC-I73T expression causes lumen loss. (A) Immunofluorescence microscopy of EEA1 and proSFTPC in SFTPC-WT/WT and WT/I73T fdAT2. Arrowhead, example enlarged SFTPC-negative endosome. Scale bar, 10µm. (B) EEA1 positive vesicle volume (inferred from diameter) in n=3 biologically independent SFTPC-WT/WT and WT/I73T fdAT2 lines. Mann-Whitney test. (C) Immunofluorescence microscopy of M6PR and proSFTPC localisation in SFTPC-WT/WT and WT/I73T fdAT2. Scale bar, 10µm. (D) Lysotracker staining and lysosome volume (inferred from area) in n=3 biologically independent SFTPC-WT/WT and WT/I73T fdAT2 lines. Mann-Whitney test. Scale bar, 10µm. (E) Immunofluorescence microscopy of EEA1 and proSFTPC localisation in SFTPC-WT/WT and WT/I73T fdAT2 induced for 0-10 days. Scale bar, 10µm. (F) Percentage of organoids with lumen present when expressing either HA-SFTPC-WT or HA-SFTPC-I73T for 0-10 days. n=50 for each for each of n=3 biologically independent lines. One way ANOVA with Sidak’s multiple comparisons. (G) Fold change in early endosome volume in fdAT2 expressing HA-SFTPC-WT or HA-SFTPC-I73T for 0-10 days. Kruskal-Wallis test with Dunn comparison. ****=p<0.0001, ***=p<0.001, *=p<0.05, ns = not significant.

Early endosomes undergo a regulated maturation process to become late endosomes through acquisition of acid hydrolases and a resulting decrease in pH^38^. We therefore hypothesised that the altered morphology of these early compartments may affect the formation and function of later compartments. Consistent with this, the late endosome marker M6PR, typically localised to intracellular puncta as seen in control organoids, was also redistributed in SFTPC-I73T-expressing organoids (Fig. 3C). Because the defect in SFTPC-I73T maturation is at the level of the MVB, we predicted that lysosomal compartments may be unaffected by SFTPC-I73T expression. Positive lysotracker® staining confirmed the presence of acidic vesicles which were not enlarged in SFTPC-I73T-expressing organoids, indicating that the observed endosomal perturbation does not prevent acidification associated with endosomal maturation or lysosomal size (Fig. 3D), consistent with findings in PSC-iAT2 and mouse models^13,14,16^. Together this suggests the observed phenotype arises from impaired trafficking at enlarged early endosomes, which serve as critical sorting platforms in polarised epithelial cells.

To define a causal relationship between loss of polarity and endosomal dysfunction, we investigated the changes in endosome volume and lumenogenesis in an inducible HA-tagged SFTPC-WT or I73T-expressing organoids. These lines were derived by transducing fdAT2 with a doxycycline-inducible lentiviral vector harbouring HA-tagged SFTPC and EF1a promoter-expressing tagRFP (suppl Fig. 3A&B). Transduced cells were sorted for RFP-positivity, and expression of HA-tagged SFTPC was induced for 10 days, after which those expressing SFTPC-I73T phenocopied their base edited counterparts (suppl Fig. 3C&D).

We used these organoids to undertake a time course (Fig. 3E-G); organoids induced to express HA-SFTPC-WT grew with defined lumens and intracellular punctate staining of SFTPC which did not co-localise with EEA1, as expected (Fig. 3E). After just 2 days of HA-SFTPC-I73T expression there was a significant increase in early endosome volume, but redistribution of SFTPC to the plasma membrane lagged behind (Fig. 3F&G), suggesting that altered endosome morphology precedes SFTPC redistribution. At later timepoints, there was more dramatic cell surface SFTPC accumulation and EEA1-positive endosomes continued to increase in volume. This time-dependent phenotype confirms a causal relationship between SFTPC-I73T expression and endosomal dysfunction and supports previous observations *ex-* and *in vivo*^14,16^.

The observed loss of lumens caused by SFTPC-I73T expression occurred early in the time course, suggesting that aberrant accumulation of SFTPC intermediates in early compartments is sufficient to disrupt lumenogenesis (Fig. 3E&F). To deconvolute whether enlarged EEA1-positive endosomes were a specific consequence of this SFTPC variant or reflect a result of endosomal protein accumulation, we measured early endosomal compartments in fdAT2 and mouse models deficient of the E3 ligase ITCH, which we have previously demonstrated is required for trafficking of SFTPC to late compartments^17^. SFTPC co-localised with EEA1 in enlarged structures in both ITCH-deplete fdAT2 (Fig. 4A-C) and ITCH-null mouse AT2 (Fig. 4D-E) suggesting the accumulation of immature SFTPC is sufficient to drive altered endosome morphology.

**Fig 4.**
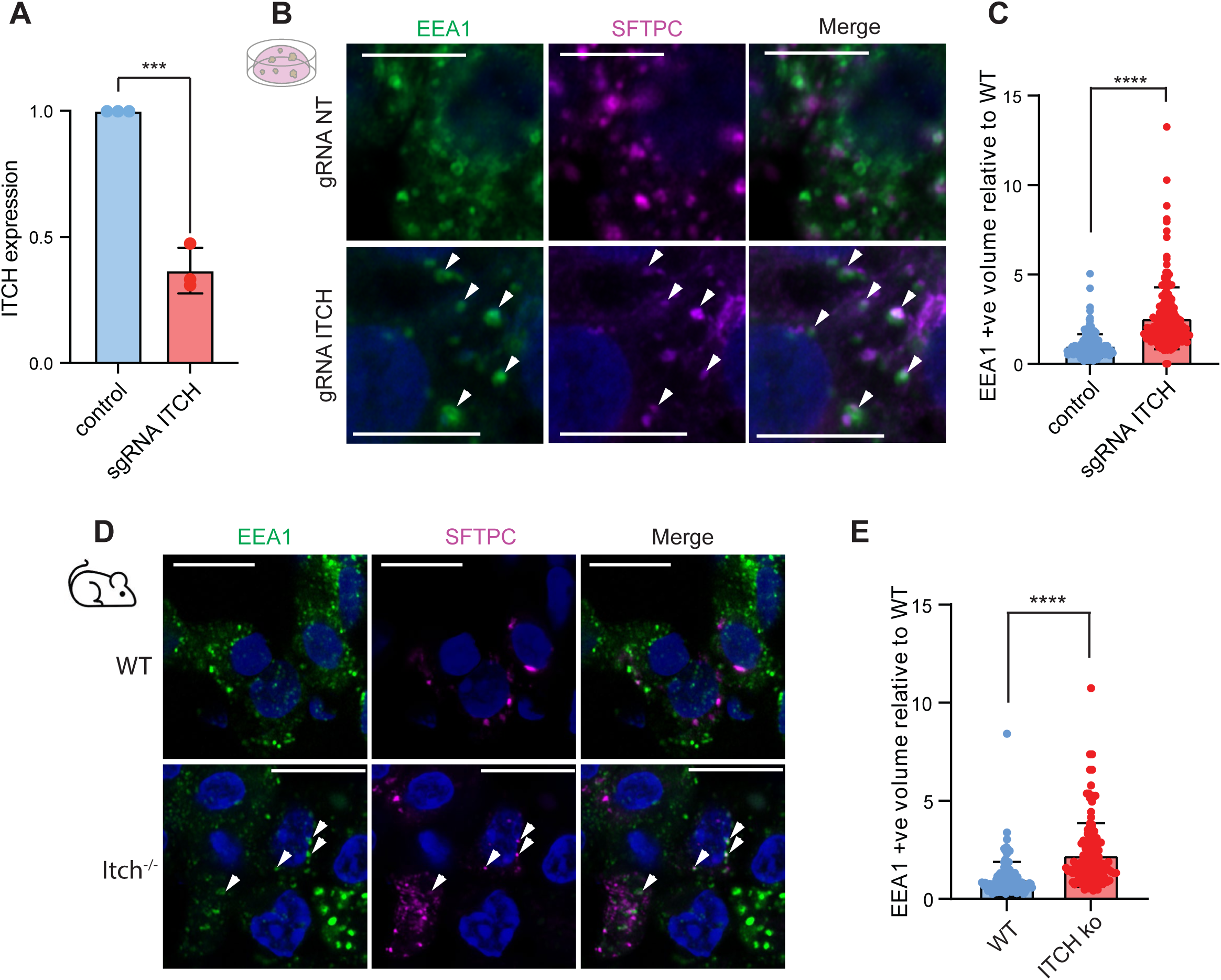
Disruption of SFTPC maturation via ITCH depletion also causes endosomal enlargement. (A) ITCH expression by qPCR in control fdAT2 and those depleted of ITCH using inducible CRISPR interference^17^. Expression relative to GAPDH and normalised to controls. Unpaired t-test of n=3 biologically independent fdAT2 lines. (B) Immunofluorescence microscopy of EEA1 and proSFTPC in control and ITCH depleted fdAT2. Arrowheads, example SFTPC-containing EEA1 positive vesicles. Scale bar, 5µm. (C) Early endosome volume in control and ITCH deplete fdAT2 from n=3 biologically independent lines. Mann Whitney test. (D) Immunofluorescence microscopy of EEA1 and proSFTPC in control and ITCH null mice. Arrowheads, example SFTPC-containing EEA1 positive vesicles. Scale bar, 10µm. (E) Early endosome volume in control and ITCH null mouse AT2. Mann Whitney test. ****=p<0.0001.

Though our organoid model provides valuable mechanistic insights, it does not reproduce the *in vivo* environment, where AT2 cells grow at an air-liquid interface. To determine whether the observed phenotypes extend beyond the organoid system, we cultured fdAT2 cells as a monolayer by dissociating organoids and plating them at high density on transwell inserts (Fig. 5A). After 48 hours of culture, fdAT2 were kept submerged or air-exposed by removal of apical media. We confirmed that parental fdAT2 form a confluent monolayer that maintains SFTPC expression for at least 10 days after air exposure (Fig. 5B&C), unlike primary AT2 which readily dedifferentiate^39–43^. We next investigated the impact of SFTPC-I73T expression in this system. Both control and I73T-fdAT2 formed a confluent monolayer 48 hours after plating; whilst control cells maintained this monolayer, air-exposure of I73T-fdAT2 resulted in the appearance of large gaps in the cell layer within 48 hours (Fig. 5D and suppl Fig. 4). 7-AAD staining revealed this was not due to excess cell death in I73T-fd2 over its control counterpart when placed at ALI, suggesting a loss of epithelial integrity rather than altered viability (Fig. 5E&F). Notably, remote from the areas of epithelial integrity loss, air-exposed I73T-fdAT2 grew in a thicker layer with phenotypic similarity to the clusters of hyperplastic AT2 cells seen in patients^13,22,44^(Fig. 5G&H). Z-stack immunofluorescence microscopy of harvested transwell inserts revealed intracellular punctate staining of SFTPC in control cells, whilst intracellular accumulation and redistribution of SFTPC to the cell surface was observed in both submerged and air-exposed I73T-fdAT2 (Fig. 5G).

**Fig 5.**
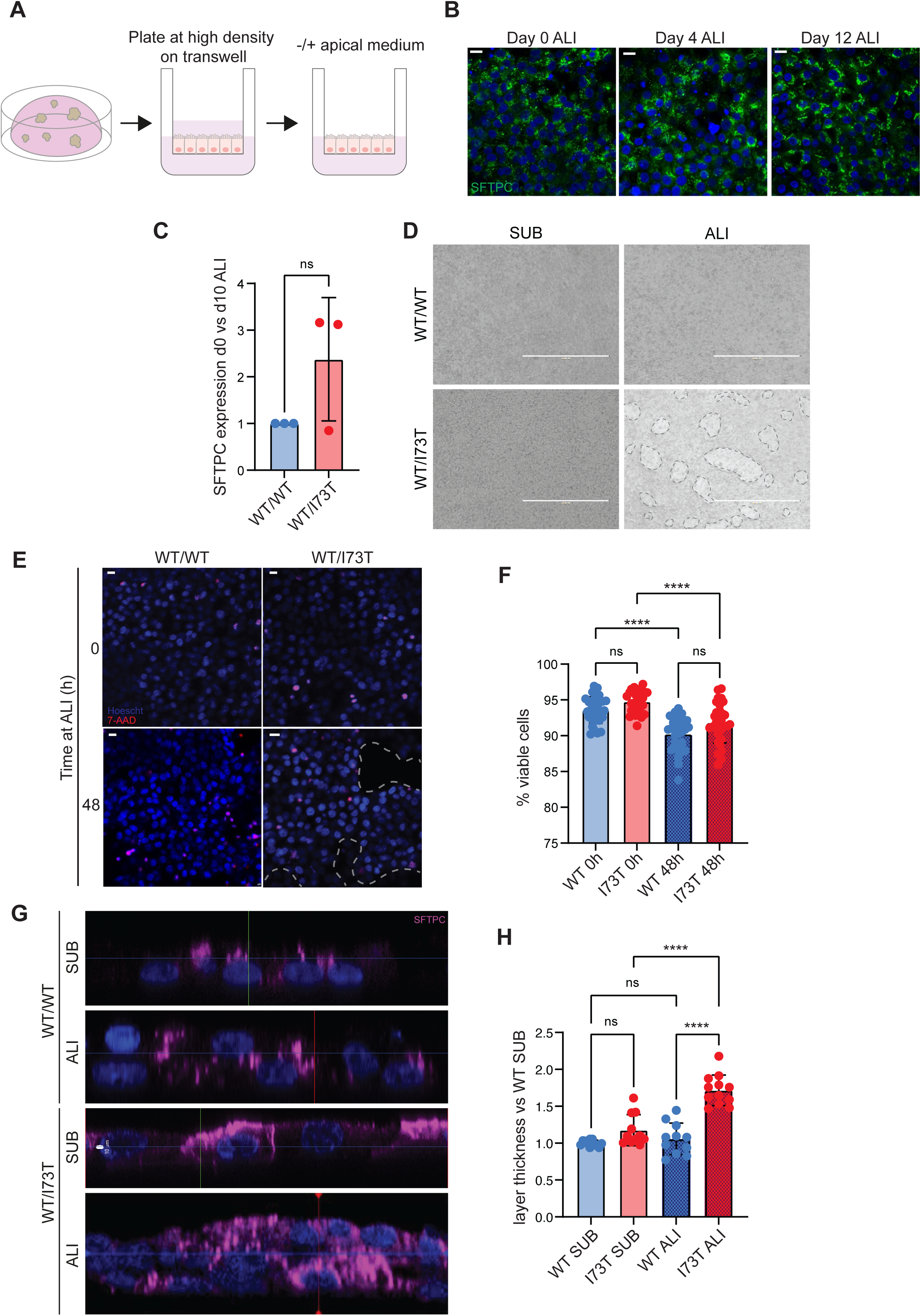
FdAT2 expressing SFTPC-I73T fail to maintain epithelial integrity when grown at air-liquid interface. (A) Schematic of plating fdAT2 for air-liquid interface (ALI) culture. (B) Immunofluorescence microscopy of proSFTPC in fdAT2 grown at ALI for up to 12 days. Scale bar, 10µm. (C) SFTPC expression by qPCR in n=3 biologically independent fdAT2 lines at d10 vs d0 ALI culture. Expression relative to GAPDH and normalised to d0. One sample Wilcoxon t-test. (D) Brightfield imaging of fdAT2 monolayer; discontinuities demarcated with dotted lines. Scale bar, 1000µm. (E) Confocal imaging of fd2 monolayer stained with Hoescht and 7-AAD, with viability of n=3 biologically independent fdAT2 lines quantified in (F) using one way ANOVA. (G) Immunofluorescence z-stack imaging of SFTPC localisation grown submerged (SUB) or at ALI; cell layer depth in n=3 biologically independent fdAT2 lines quantified in (H) using a Welch ANOVA test. ****=P<0.0001, ns = not significant.

In a manner phenocopying 3D culture, the redistribution of SFTPC-I73T in 2D culture was associated with enlargement of early endosomes but not acidic, later compartments (Fig. 6A-C). In keeping with this, lamellar body biogenesis appeared unaffected, with no observable changes in the size or distribution of LBs in I73T-fdAT2 (Fig. 6D). Interestingly, air exposure further exacerbated the endosomal defects with a two-fold increase in early endosome volume (Fig. 6A&B), suggesting that perturbations in early endosome size and function may be driving the observed loss of epithelial integrity. Air exposure was also sufficient to cause enlargement of EEA1-positive endosomes in air-exposed control fdAT2, likely reflecting the increased trafficking demands associated with AT2 maturation and surfactant secretion upon air exposure^45–48^(Fig. 6B).

**Fig 6.**
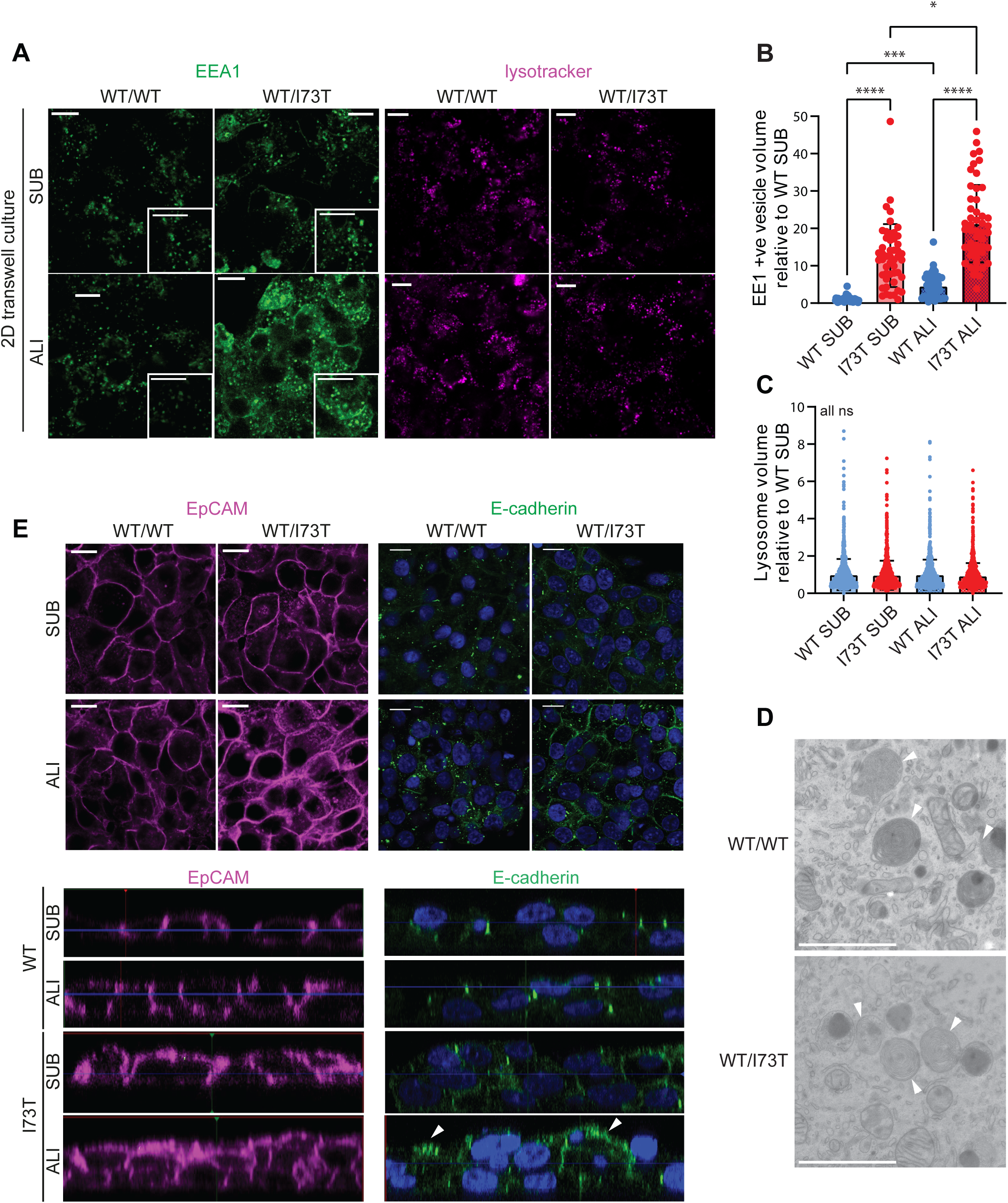
Air-liquid interface culture exacerbates the endosomal phenotype caused by SFTPC-I73T expression. (A) Immunofluorescence imaging (left panel) / lysotracker imaging (right panel) of fdAT2 grown submerged or at ALI for 48 hours. Scale bar, 10µm. (B&C) Quantification of early endosome and lysosome volumes in n=3 biologically independent fdAT2 lines. Kruskal Wallis test with Dunn’s multiple comparison. (E) Immunofluorescence microscopy of EpCAM and E-Cadherin in fdAT2 grown submerged or at ALI for 48 hours (top panels, confocal slice; bottom panels, z-stacks). Scale bar, 10µm. (D) Transmission electron microscopy imaging of SFTPC-WT/WT and WT/I73T fdAT2. Arrowheads, lamellar bodies. Scale bar, 200nm. ****=p<0.0001, ***=p<0.001, *=p<0.05, ns=not significant.

Disrupted endosomal morphology in I73T-fdAT2 was accompanied by defects in the localisation of cargo sorting and cell adhesion complexes, with a striking redistribution of both EpCAM and E-cadherin to the apical plasma membrane (Fig. 6E). This mislocalisation suggests apicobasal polarity remains perturbed even in a system where distinct apical and basal cues are present, reinforcing that this phenotype is not an artefact of three-dimensional culture. Indeed, the excessive plasma membrane accumulation of these proteins in air-exposed cells may impair the function of cell-cell junctions and explain the loss of epithelial integrity observed under these conditions.

Given the striking disruption to epithelial barrier integrity in I73T-fdAT2, we wished to investigate whether these defects had an impact on epithelial repair, the dysregulation of which is a hallmark of IPF. In two lines, expression of the SFTPC-I73T variant resulted in such a strong loss-of-integrity phenotype that this was impossible; however we discovered that when cells were plated to excess (approximately double the standard plating density, suppl Fig. 5), we were able to overcome loss of epithelial integrity sufficiently to undertake wound healing assays in line 2 (Fig. 7A). This variability in phenotype likely in part reflects *in vivo* differences in disease penetrance and severity. Expression of SFTPC-I73T slowed wound healing under both submerged and air-exposed culture conditions, though impairment of healing was exacerbated by air exposure. This was evident both from absolute measures of wound healing (Fig. 7A&B) and cross-sectional assays of light penetration, which confirmed that healing was less complete in I73T-fdAT2 (Fig. 7C).

**Fig 7.**
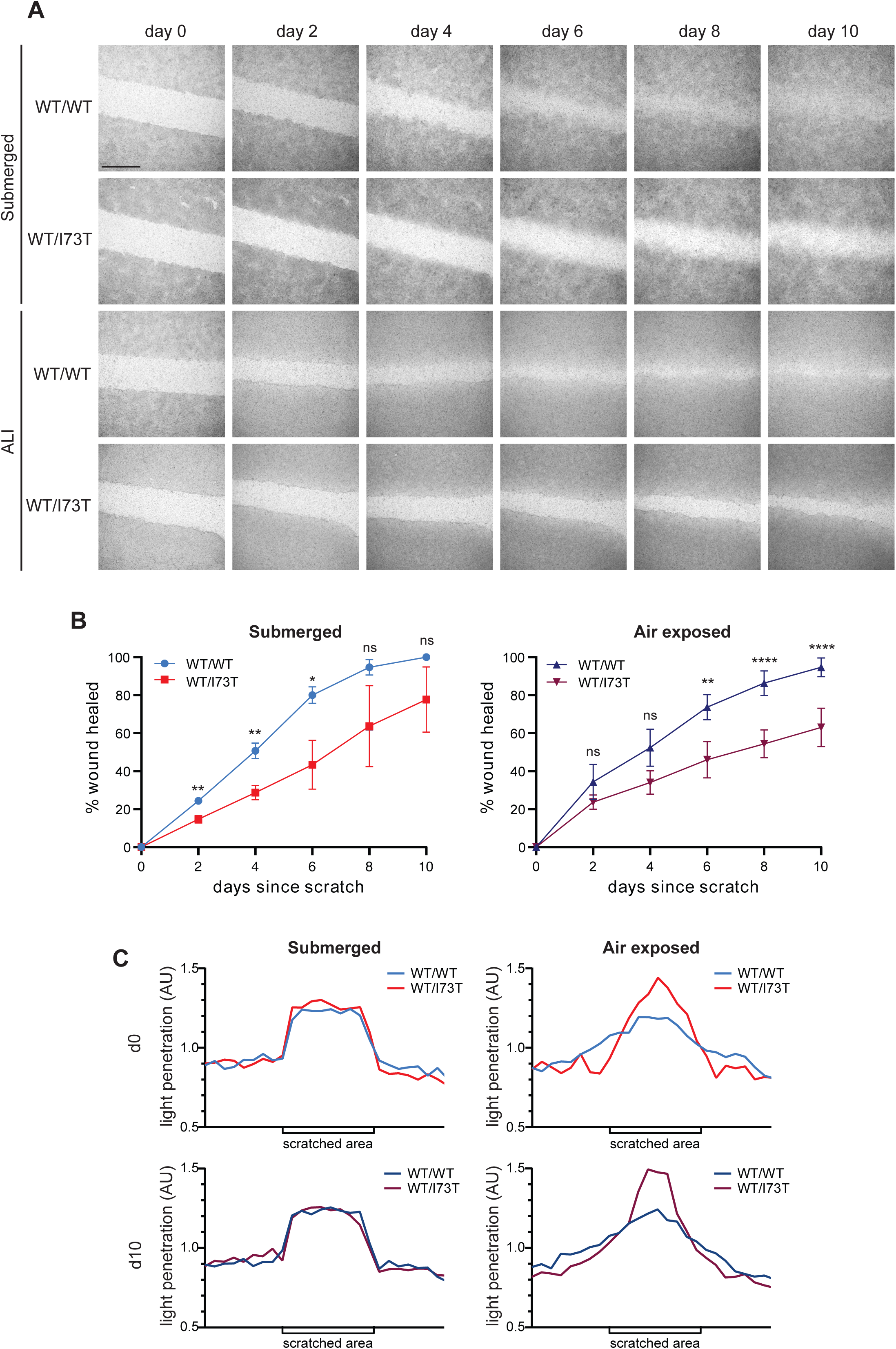
SFTPC-I73T expression impairs wound healing. (A) Sequential imaging of wound healing in SFTPC-WT/WT or WT/I73T grown submerged or at ALI. Scale bar, 500µm. (B) Percentage wound healed with time in submerged culture (left panel) and at ALI (right panel). Two-way ANOVA of n=3 independent repeats using line 2. (C) Representative light penetration across imaging field at d0 and d10 post wounding. ****=p<0.0001, **=p<0.005, *=p<0.05, ns=not significant.

Taken together, using endogenous gene editing of fdAT2, we herein demonstrate the consequences of SFTPC-I73T expression on endosomal function, polarity, and the abundance of endosomally trafficked cargoes at the plasma membrane. Together this results in the impairment of epithelial integrity and repair, providing further insight into the pathogenic mechanisms of SFTPC-I73T expression.

## Discussion

In this study, we aimed to create an endogenous disease model to further understand the mechanisms by which SFTPC-I73T expression induces alveolar epithelial cell dysfunction in familial pulmonary fibrosis. Using base editing of fdAT2, which are quicker to derive and more readily genetically manipulable alternative to iAT2, we developed a heterozygous expression model of SFTPC-I73T. We then established fdAT2 cultures at ALI, allowing us for the first time to model the phenotypic changes in SFTPC-I73T-expressing fdAT2 driven by air-exposure. This study reports the first successful application of base-editing to differentiated human alveolar epithelial cells, highlighting its utility not only for generating physiologically relevant preclinical models but also as a possible strategy for therapeutic correction of disease-causing mutations *in vivo*. In our study, bystander editing was not functionally important but does highlight a limitation of this approach.

Our model phenocopied the intracellular accumulation and aberrant redistribution of SFTPC-I73T previously described in immortalised cell lines, iAT2, mouse models, and patient tissue^12,14,16^. Mechanistic studies of SFTPC-I73T trafficking have shown that failure of ubiquitination results in aberrant accumulation of SFTPC intermediates in early endosomal compartments which are then recycled to the plasma membrane^15–17,22^. This results in impaired autophagic flux and mitophagy in AT2, although it remains mechanistically unclear exactly how SFTPC accumulation causes these perturbations and initiates the fibrotic cascade^13,14,16^. Here we demonstrate that SFTPC-I73T expression in fdAT2 organoids drives time-dependent endosomal dysfunction and loss of spatial organisation. The accumulation of SFTPC-I73T intermediates leads to the enlargement of early endosomes, which constitute major sorting platforms in polarised epithelial cells^37,49–51^. This is followed by redistribution of SFTPC-I73T, key apical and basal cargoes, and cell adhesion proteins. This early endosomal enlargement and impaired endocytic sorting mirrors defects seen in other diseases of endosomal trafficking abnormalities, including Niemann-Pick type C and Alzheimer’s disease^52,53^.

Intact cell-cell junctions are required for epithelial integrity and for the establishment and maintenance of apico-basal polarity^54^. In our inducible system, SFTPC-I73T redistribution precedes the loss of organisation in I73T-fdAT2, suggesting defective endosomal sorting is an early, time-dependent driver of polarity loss. Our observed phenotype is in keeping with observations in iAT2 models^16^ and the ultrastructural abnormalities reported in embryonic mouse lungs carrying the same variant^14^. Importantly, loss of polarity was still apparent in 2-dimensional culture where clear apical and basal cues are present, indicating this is not an artefact of 3D culture.

Loss of epithelial polarity is seen in pulmonary fibrosis. Bleomycin-injured primary AT2 cells lose apico-basal polarity^55^ and aberrant basaloid cells, which are IPF-specific aberrant epithelial-derived cells^56,57^, show marked transcriptomic upregulation of mesenchymal signatures, characterised by loss of cell-junctions and polarity. Our data raise the possibility that endosomal dysfunction may be one trigger of epithelial polarity loss and dysfunction. Similar endolysosomal and membrane trafficking defects are seen in Hermansky-Pudlak syndrome (HPS), a rare autosomal recessive hereditary disease which causes FPF; HPS organoid models also display marked structural abnormalities and mesenchymal cell emergence^58–60^. The very nature of organoid culture essentially reproduces a serial injury model, so our and others’ observed phenotypes may in fact phenocopy the consequences of the types of “second hits” that *in vivo* trigger disease. Whether prolonged SFTPC-I73T expression results in any fundamental change in AT2 phenotype, perhaps towards the type of aberrant intermediate cells seen in IPF, is challenging to determine in the culture conditions used for iAT2 and fdAT2 which are designed to maintain a stable AT2 phenotype but co-culture models or PCLS cultures which require less prescriptive culture media may be able to address these questions.

In our I73T-fdAT2, mislocalisation and accumulation of EpCAM and E-cadherin were associated with epithelial disorganisation and impaired wound closure, consistent with disruption of their roles in maintaining cohesion and supporting collective migration^61,62^. This phenotype became more profound upon air-exposure, highlighting the increased vulnerability of SFTPC-I73T-expressing epithelium under more physiologically relevant conditions. In many of those with an SFTPC-I73T variant, disease presents in the neonatal period^63^, likely reflecting AT2 decompensation upon air exposure at birth. Cell-cell adhesion and tight junction defects are a recognised feature of pulmonary fibrosis more broadly^64–67^ and bleomycin injury to alveolar epithelial cells in mice significantly disrupts claudin expression, which contributes to epithelial barrier dysfunction^68^. Our findings support and extend these observations by identifying endosomal mis-sorting of adhesion proteins as a potential upstream driver of junctional instability and repair failure.

Our inducible SFTPC-I73T system has allowed us to capture upstream events that precede the autophagy defects reported in iAT2 and mouse disease models. Several core components of the endosomal machinery, including Rab35, the ESCRT complex and AP-2, not only mediate cargo sorting but also play essential roles in autophagosome formation and maturation^69,70^. Early endosomes have been implicated in autophagy and mitophagy regulation^71^, whilst recent studies have shown that autophagy machinery can directly target damaged early endosomes for degradation^72^, positioning them as both regulators and substrates of the pathway. These mechanistic intersections suggest that early endosomal dysfunction may underlie downstream autophagy perturbations and are worthy of further study. Similar convergence of endosomal and autophagic stress has been extensively described in several neurodegenerative diseases, including Alzheimer’s and Huntington’s diseases, further underscoring the relevance of this axis in chronic disease^73,74^. Together, these findings position endosomal disruption as a potential initiating event in SFTPC-I73T–driven epithelial dysfunction and highlight the value of complementary models for resolving the temporal progression of disease.

We undertook all experiments at early passage to focus on understanding the functional consequences of SFTPC-I73T redistribution, but observed significant phenotypic drift in one biological line at later passage, despite maintained genotypic stability. This was marked by a loss of surface SFTPC expression and may represent a compensatory response to chronic mutant protein accumulation, though plausibly could represent the consequences of off-target editing in this line. It may also indicate a transition toward an altered cell state that is incompletely facilitated under defined culture conditions. These changes underscore the importance of monitoring phenotypic stability over time in any endogenous disease model. The variation in severity of phenotype between the three biological lines, particularly when assessing wound healing, implicates other intrinsic factors in determining the degree of AT2 dysfunction and likely reflects the clinical heterogeneity seen in patients.

Together, these findings define a mechanism by which SFTPC-I73T drives progressive AT2 dysfunction through endosomal disruption, altered endocytic sorting, and loss of polarity. By modelling SFTPC-I73T expression at endogenous levels and at an air–liquid interface, we capture key features of disease pathogenesis and demonstrate how air exposure exacerbates epithelial vulnerability. The use of base editing in differentiated human alveolar cells establishes an important platform for both mechanistic studies and therapeutic development. More broadly, this work implicates polarity loss and epithelial breakdown as unifying features across familial and sporadic pulmonary fibrosis.

## Acknowledgement and funding

ENR and JAD are funded by an MRC Clinician Scientist Fellowship (MR/S005552/1). JAD is also funded by an Action for Pulmonary Fibrosis Mike Bray Fellowship (MBF2023_001). KL is supported by the Korea University Grant and by the National Research Foundation of Korea (NRF) grant funded by Korea government (MSIT) (RS-2024-00342036 and RS-2023-00221182). JRE is supported by a Sir Henry Dale Fellowship jointly funded by the Wellcome Trust and the Royal Society (216370/Z/19/Z). LEM was supported in part by the National Institute of General Medical Sciences of the National Institutes of Health under Award Number P20GM103499. SJM is supported by the MRC (MCMB MR/V028669/1 and MR/Y011813/1), EPSRC (EP/R03558X/1), Cambridge Biomedical Research Centre (BRC-1215-20014); Asthma+Lung UK (ALUK), and the Victor Philip Dahdaleh Foundation. ELR is supported by the Medical Research Council (MR/P009581/1) and Wellcome Discovery Award (225221/Z/22/Z). The authors would also like to acknowledge the support of the CIMR Flow Cytometry Core Facility.

## Author contributions

**Eimear Rutherford:** Conceptulisation, methodology, investigation, analysis, writing - original draft. **Dawei Sun:** Conceptulisation, methodology, investigation, analysis, writing - review and editing. **Kyungtae Lim:** Conceptulisation, methodology, investigation, analysis, writing - review and editing **James Edgar:** Methodology, investigation, writing - review and editing. **Lydia Matesic:** Resources, methodology, writing - review and editing. **Stefan Marciniak:** Conceptualisation, resources, supervision, writing - review and editing**. Emma Rawlins:** Conceptualisation, supervision, funding acquisition, writing - review and editing. **Jennifer Dickens:** conceptualisation, supervision, validation, investigation, methodology, project administration, writing - original draft.

## Materials and Methods

Details of tissue and cell sources, antibodies, expression plasmids, reagents, equipment and software are included in table 1.

**Table 1:**
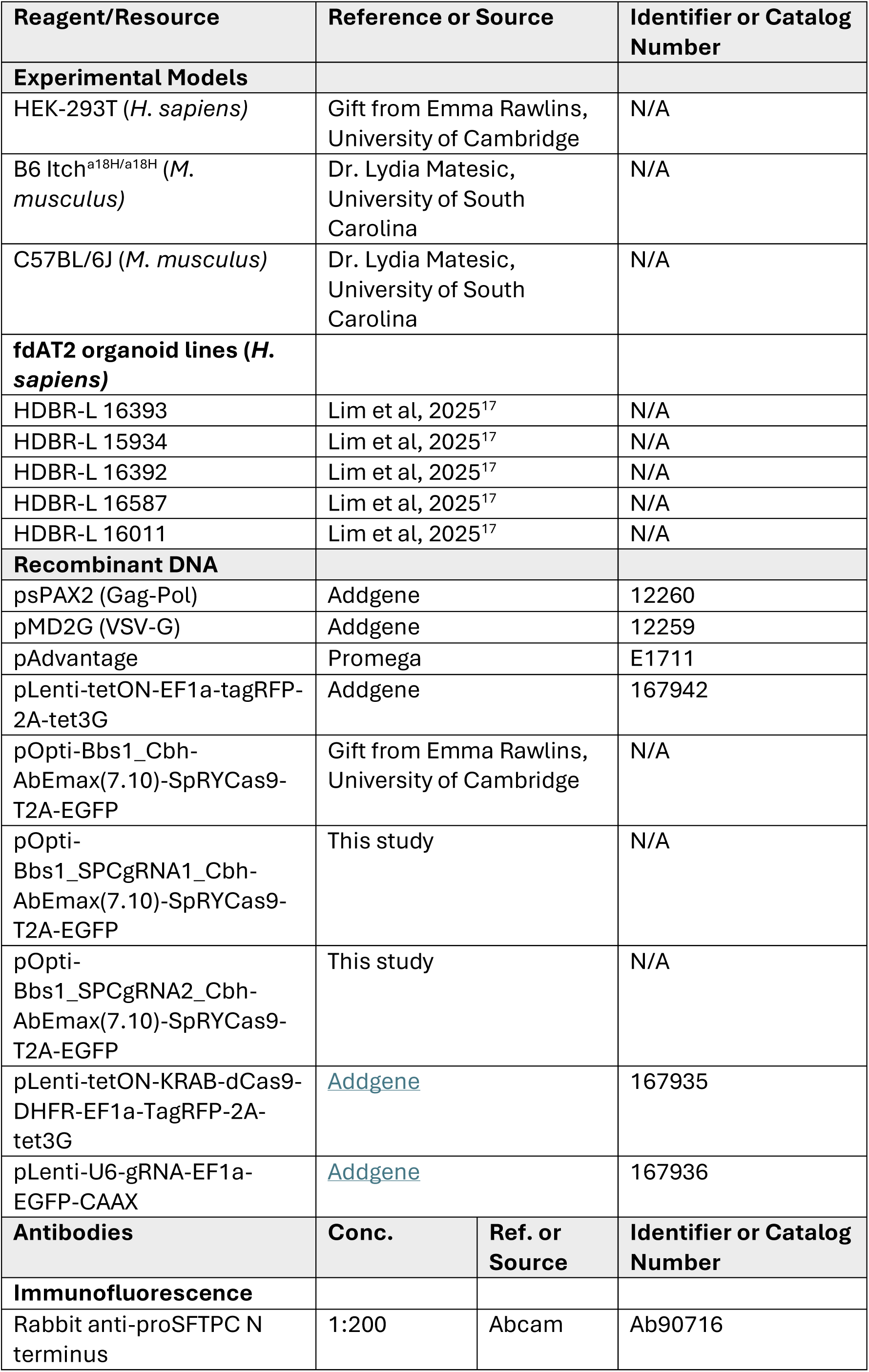
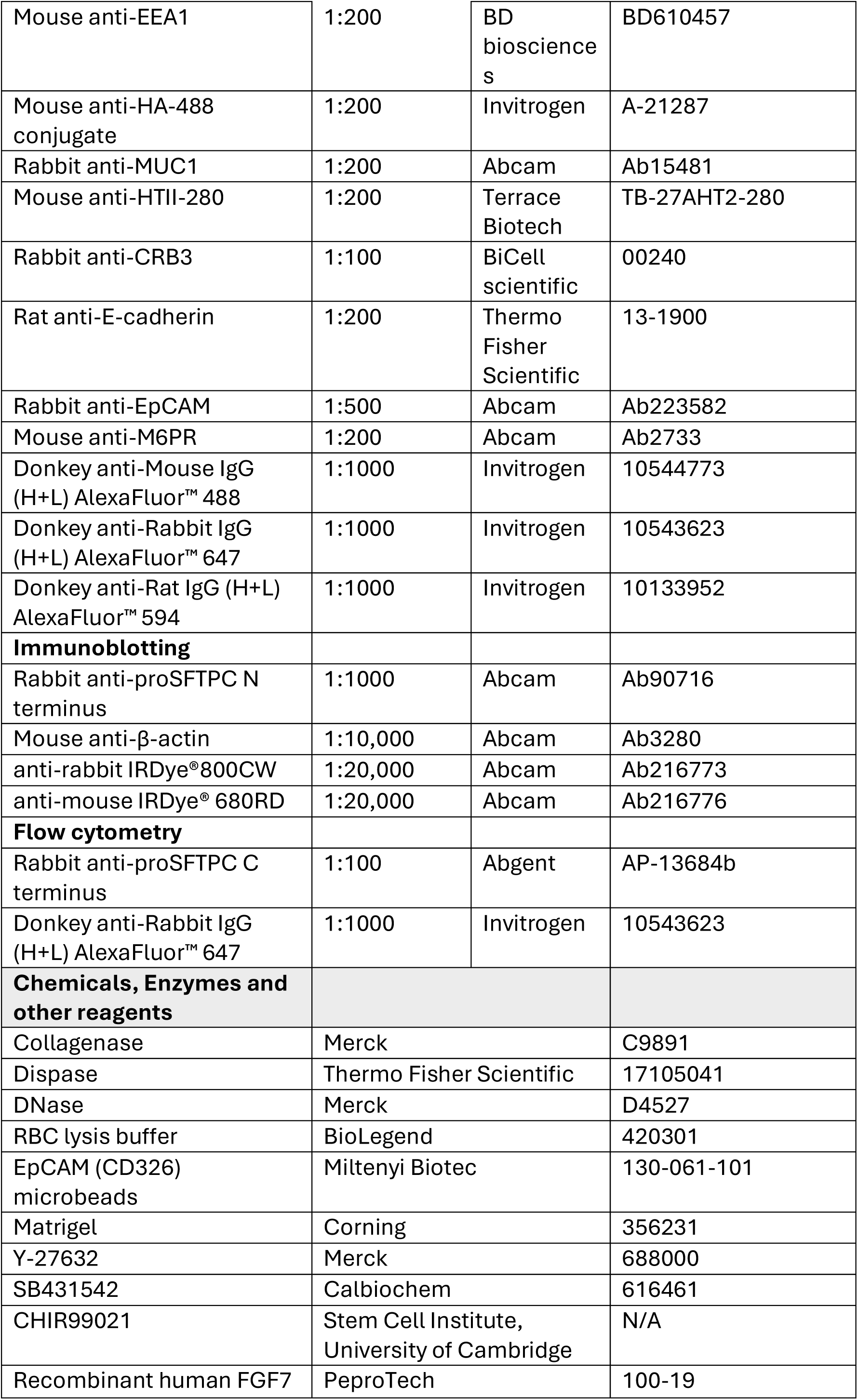
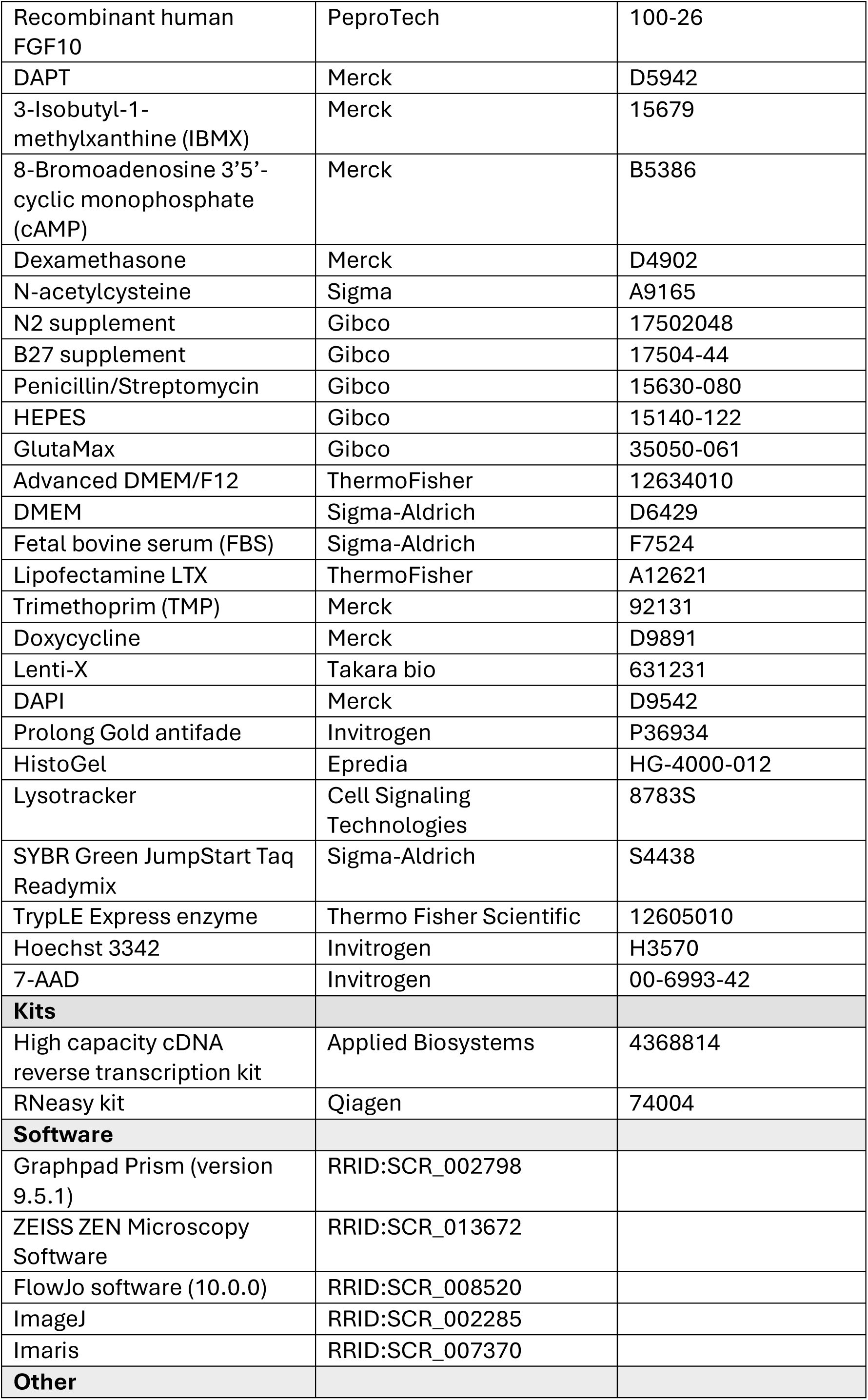
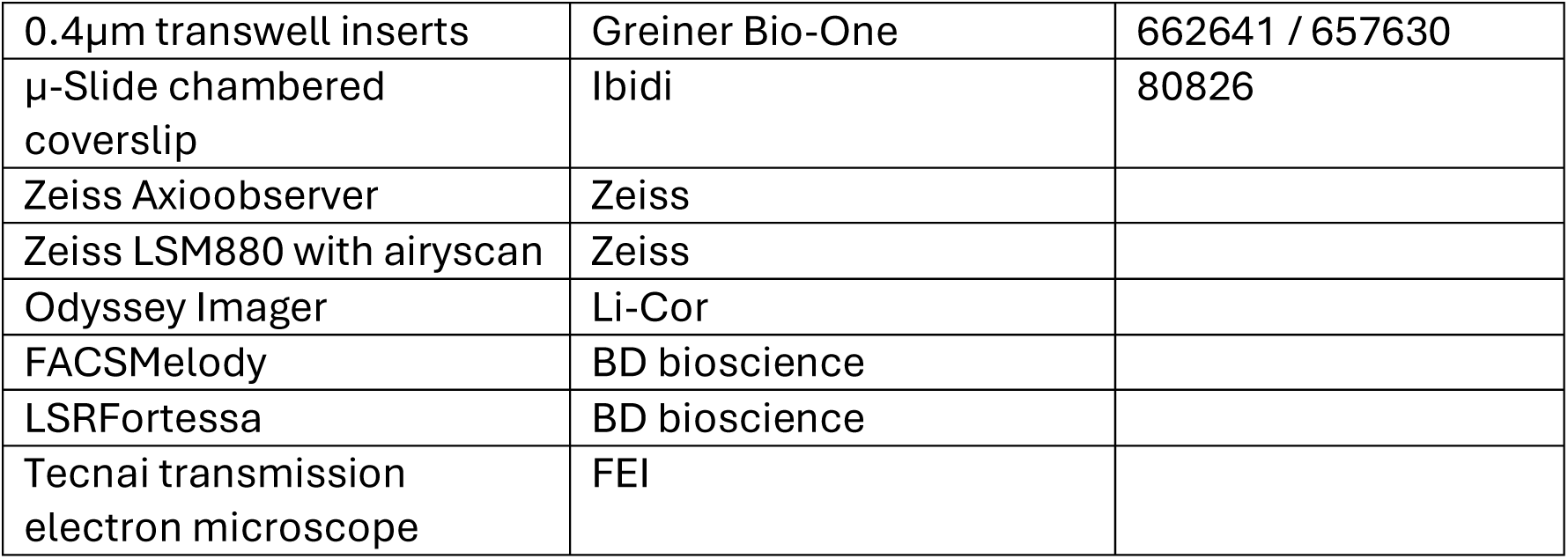
Reagents and Tools.

| Reagent/Resource | Reference or Source |  | Identifier or Catalog Number |
| --- | --- | --- | --- |
| Experimental Models |  |  |  |
| HEK-293T (*H. sapiens*) | Gift from Emma Rawlins, University of Cambridge |  | N/A |
| B6 Itch<sup>a18H/a18H</sup> (*M. musculus*) | Dr. Lydia Matesic, University of South Carolina |  | N/A |
| C57BL/6J (*M. musculus*) | Dr. Lydia Matesic, University of South Carolina |  | N/A |
| fdAT2 organoid lines (*H. sapiens*) |  |  |  |
| HDBR-L 16393 | Lim et al, 2025<sup>17</sup> |  | N/A |
| HDBR-L 15934 | Lim et al, 2025<sup>17</sup> |  | N/A |
| HDBR-L 16392 | Lim et al, 2025<sup>17</sup> |  | N/A |
| HDBR-L 16587 | Lim et al, 2025<sup>17</sup> |  | N/A |
| HDBR-L 16011 | Lim et al, 2025<sup>17</sup> |  | N/A |
| Recombinant DNA |  |  |  |
| psPAX2 (Gag-Pol) | Addgene |  | 12260 |
| pMD2G (VSV-G) | Addgene |  | 12259 |
| pAdvantage | Promega |  | E1711 |
| pLenti-tetON-EF1a-tagRFP-2A-tet3G | Addgene |  | 167942 |
| pOpti-Bbs1_Cbh-AbEmax(7.10)-SpRYCas9-T2A-EGFP | Gift from Emma Rawlins, University of Cambridge |  | N/A |
| pOpti-Bbs1_SPCgRNA1_Cbh-AbEmax(7.10)-SpRYCas9-T2A-EGFP | This study |  | N/A |
| pOpti-Bbs1_SPCgRNA2_Cbh-AbEmax(7.10)-SpRYCas9-T2A-EGFP | This study |  | N/A |
| pLenti-tetON-KRAB-dCas9-DHFR-EF1a-TagRFP-2A-tet3G | Addgene |  | 167935 |
| pLenti-U6-gRNA-EF1a-EGFP-CAAX | Addgene |  | 167936 |
| Antibodies | Conc. | Ref. or Source | Identifier or Catalog Number |
| Immunofluorescence |  |  |  |
| Rabbit anti-proSFTPC N terminus | 1:200 | Abcam | Ab90716 |
| Mouse anti-EEA1 | 1:200 | BD biosciences | BD610457 |
| Mouse anti-HA-488 conjugate | 1:200 | Invitrogen | A-21287 |
| Rabbit anti-MUC1 | 1:200 | Abcam | Ab15481 |
| Mouse anti-HTII-280 | 1:200 | Terrace Biotech | TB-27AHT2-280 |
| Rabbit anti-CRB3 | 1:100 | BiCell scientific | 00240 |
| Rat anti-E-cadherin | 1:200 | Thermo Fisher Scientific | 13-1900 |
| Rabbit anti-EpCAM | 1:500 | Abcam | Ab223582 |
| Mouse anti-M6PR | 1:200 | Abcam | Ab2733 |
| Donkey anti-Mouse IgG (H+L) AlexaFluor™ 488 | 1:1000 | Invitrogen | 10544773 |
| Donkey anti-Rabbit IgG (H+L) AlexaFluor™ 647 | 1:1000 | Invitrogen | 10543623 |
| Donkey anti-Rat IgG (H+L) AlexaFluor™ 594 | 1:1000 | Invitrogen | 10133952 |
| **Immunoblotting** |  |  |  |
| Rabbit anti-proSFTPC N terminus | 1:1000 | Abcam | Ab90716 |
| Mouse anti-β-actin | 1:10,000 | Abcam | Ab3280 |
| anti-rabbit IRDye®800CW | 1:20,000 | Abcam | Ab216773 |
| anti-mouse IRDye® 680RD | 1:20,000 | Abcam | Ab216776 |
| **Flow cytometry** |  |  |  |
| Rabbit anti-proSFTPC C terminus | 1:100 | Abgent | AP-13684b |
| Donkey anti-Rabbit IgG (H+L) AlexaFluor™ 647 | 1:1000 | Invitrogen | 10543623 |
| **Chemicals, Enzymes and other reagents** |  |  |  |
| Collagenase | Merck |  | C9891 |
| Dispase | Thermo Fisher Scientific |  | 17105041 |
| DNase | Merck |  | D4527 |
| RBC lysis buffer | BioLegend |  | 420301 |
| EpCAM (CD326) microbeads | Miltenyi Biotec |  | 130-061-101 |
| Matrigel | Corning |  | 356231 |
| Y-27632 | Merck |  | 688000 |
| SB431542 | Calbiochem |  | 616461 |
| CHIR99021 | Stem Cell Institute, University of Cambridge |  | N/A |
| Recombinant human FGF7 | PeproTech |  | 100-19 |

|  |  |  |
| --- | --- | --- |
| Recombinant human FGF10 | PeproTech | 100-26 |
| DAPT | Merck | D5942 |
| 3-Isobutyl-1-methylxanthine (IBMX) | Merck | 15679 |
| 8-Bromoadenosine 3'5'-cyclic monophosphate (cAMP) | Merck | B5386 |
| Dexamethasone | Merck | D4902 |
| N-acetylcysteine | Sigma | A9165 |
| N2 supplement | Gibco | 17502048 |
| B27 supplement | Gibco | 17504-44 |
| Penicillin/Streptomycin | Gibco | 15630-080 |
| HEPES | Gibco | 15140-122 |
| GlutaMax | Gibco | 35050-061 |
| Advanced DMEM/F12 | ThermoFisher | 12634010 |
| DMEM | Sigma-Aldrich | D6429 |
| Fetal bovine serum (FBS) | Sigma-Aldrich | F7524 |
| Lipofectamine LTX | ThermoFisher | A12621 |
| Trimethoprim (TMP) | Merck | 92131 |
| Doxycycline | Merck | D9891 |
| Lenti-X | Takara bio | 631231 |
| DAPI | Merck | D9542 |
| Prolong Gold antifade | Invitrogen | P36934 |
| HistoGel | Epredia | HG-4000-012 |
| Lysotracker | Cell Signaling Technologies | 8783S |
| SYBR Green JumpStart Taq Readymix | Sigma-Aldrich | S4438 |
| TrypLE Express enzyme | Thermo Fisher Scientific | 12605010 |
| Hoechst 3342 | Invitrogen | H3570 |
| 7-AAD | Invitrogen | 00-6993-42 |
| **Kits** |  |  |
| High capacity cDNA reverse transcription kit | Applied Biosystems | 4368814 |
| RNeasy kit | Qiagen | 74004 |
| **Software** |  |  |
| Graphpad Prism (version 9.5.1) | RRID:SCR_002798 |  |
| ZEISS ZEN Microscopy Software | RRID:SCR_013672 |  |
| FlowJo software (10.0.0) | RRID:SCR_008520 |  |
| ImageJ | RRID:SCR_002285 |  |
| Imaris | RRID:SCR_007370 |  |
| **Other** |  |  |
| 0.4µm transwell inserts | Greiner Bio-One | 662641 / 657630 |
| µ-Slide chambered coverslip | Ibidi | 80826 |
| Zeiss Axioobserver | Zeiss |  |
| Zeiss LSM880 with airyscan | Zeiss |  |
| Odyssey Imager | Li-Cor |  |
| FACSMelody | BD bioscience |  |
| LSRFortessa | BD bioscience |  |
| Tecnai transmission electron microscope | FEI |  |

### Material Availability

The human organoid lines used in the study are available from Dr. Emma L Rawlins with a completed Materials Transfer Agreement.

### Mouse tissue

Twelve-week old animals homozygous for a null allele of *Itch* (*Itch^a18H/a18H^*, B6.C3H(101)-In(2a;Itch)18H/LmatMmjax MMRRC stock #65285) have been previously described^75,76^. For this line, the *a^18H^* allele was backcrossed to C57BL/6J for 27 generations. Therefore, age- and gender-matched C57BL/6J mice were used as controls (*Itch^+/+^*) in the indicated experiments. All mice were cared for in accordance with the National Institute of Health’s Guide for the Care and Use of Laboratory Animals (8th ed., Washington, DC: 2011) and the University of South Carolina’s Institutional Animal Care and Use Committee approved all experimental protocols.

### Human fetal lung tissue

Human embryonic and fetal lung tissue was provided from terminations of pregnancy from the MRC/Wellcome Trust Human Developmental Biology Resource (London and Newcastle, University College London (UCL) site REC reference: 18/LO/0822; Project 200591; www.hdbr.org). Sample age ranged from 16-22 weeks of gestation (post-conception weeks; pcw). Sample gestation was determined by external physical appearance and measurements. Samples had no known genetic abnormalities. Sample gender was unknown at the time of collection and was not determined.

### Derivation and *in vitro* culture of fdAT2

Alveolar organoids were generated from human fetal lung tissue as previously described^17^. Briefly, lung tissue was sectioned and fragmented before dissociation to single cells using collagenase and dispase. The resulting cell suspension was filtered through a 100μm strainer and treated with RBC lysis buffer before isolation of EpCAM+ epithelial cells using EpCAM microbeads. These cells were embedded at high density in matrigel and cultured in 24-well plates in AT2 differentiation media (fdAT2 medium) for 14 days. To passage, organoids were dissociated by incubating with TrypLE at 37°C, pipetting every 2 minutes to gently break up organoid colonies. Single cells were embedded at high density in matrigel (typically 2-4 × 10^4^ cells/50µL Matrigel) and cultured in fdAT2 cell medium (Advanced DMEM/F12 supplemented with 1× GlutaMax, 1 mM HEPES and 1x penicillin/streptomycin (hereafter = “DMEM +++”), 1× B27 supplement (without Vitamin A), 1× N2 supplement, 1.25mM N-acetylcysteine, 50nM Dexamethasone, 0.1 mM 8-Bromoadenosine 3’5’-cyclic monophosphate, 0.1mM 3-Isobutyl-1-methylxanthine (IBMX), 50μM DAPT, 100 ng/ml recombinant human FGF7, 3μM CHIR99021, 10μM SB431542, and 10 μM Y-27632). Medium was replaced every 2 days and organoids were typically passaged every 7-10 days at a 1 to 10 ratio. For experiments at the air-liquid interface, organoids were dissociated and ∼1.5×10^5^ plated on 0.4µm transwell inserts coated in 20% matrigel, according to a previously published protocol^77^. After 48 hours of culture, fdAT2 were kept submerged or air-exposed by removal of apical media.

### Lentivirus production and viral transduction

The plasmid vectors used for producing lentivirus for organoid transductions are detailed in table 1. HEK293T cells were plated at a density of 2 or 8×10^5^ (for generating HA-SFTPC / base edited lines, respectively) in a 10cm dish 24 hours prior to transduction. Lipofectamine LTX was used to transfect cells with 5 or 10 µg plasmid of interest, 3 or 6µg pPAX2, 2 or 3µg pMD2G, and 2 or 3µg pAdv and media was changed the next day. Lentivirus was collected after 48 hours and filtered through a 0.45µm filter. Lentivirus generated for base editing was subsequently concentrated by ultracentrifugation at 7200 *x g* for 2 hours. Lentivirus for generating HA-SFTPC-WT or I73T lines was concentrated with Lenti-X concentrator (3:1 supernatant to Lenti-X) overnight at 4°C, spun at 1,500 *x g* for 45 min in a cold centrifuge. Pellets were resuspended in PBS to yield a 100X concentrated stock.

### Genetic manipulation of organoids

To create transduced organoid lines, single-cell dissociated fdAT2 organoids were typically infected with 25μl 100x concentrated lentivirus overnight at 37°C then embedded into matrigel and cultured in the medium for another 5-10 days before fluorescence sorting to enrich for transduced cells.

For HA-tagged SFTPC lines, HA-SFTPC-WT or I73T were cloned into pLenti-tetON-EF1a-tagRFP-2A-tet3G by Gibson assembly. Transduced cells were enriched by sorting for RFP positivity. Expression of HA-SFTPC-WT or I73T was induced by adding 2μg/ml doxycycline to fdAT2 medium.

For endogenous gene editing, the adenine base editing vector, pOpti-Bbs1_Cbh-AbEmax(7.10)-SpRYCas9-T2A-EGFP, was modified by insertion of sgRNA sequences targeting SFTPC to generate pOpti-Bbs1_SPCgRNA1_Cbh-AbEmax(7.10)-SpRYCas9-T2A-EGFP and pOpti-Bbs1_SPCgRNA2_Cbh-AbEmax(7.10)-SpRYCas9-T2A-EGFP (see table 2 for gRNA sequences). Fluorescence sorting for GFP positivity was used to enrich transduced cells. Following a period of recovery and expansion (typically 7-14 days), cells were further sorted based on cell surface SFTPC levels using C-terminal BRICHOS domain antibody staining, and editing efficiency of each population was determined by Sanger sequencing. Control lines were transduced with pOpti-Bbs1_Cbh-AbEmax(7.10)-SpRYCas9-T2A-EGFP without an sgRNA insert, hereafter called “empty vector”, and sorted for GFP positivity.

**Table 2:**
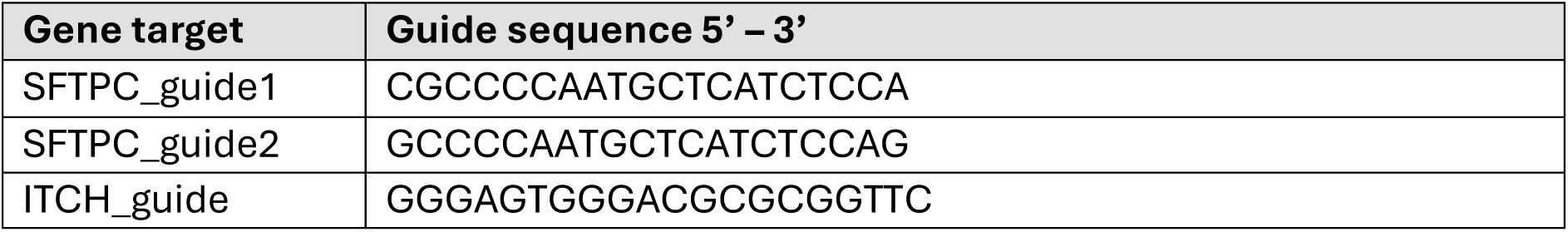
sgRNA sequences for base editing and CRISPR interference.

CRISPRi was undertaken as previously described^17^. Briefly, organoids were transduced with pLenti-tetON-KRAB-dCas9-DHFR-EF1a-TagRFP-2A-tet3G, sorted for RFP positivity then transduced with pLenti-U6-gRNA-EF1a-EGFP-CAAX containing an sgRNA targeted against *ITCH* (see table 2 for gRNA sequence). For activation of the CRISPRi system, 2μg/ml dox and 10nmol/l TMP were added to the medium for 5 days before harvesting. Three biologically independent sources were used to create n=3 HA-tagged, CRISPRi and base edited lines.

### Flow cytometry

Organoids were dissociated to single cells, washed twice in DMEM+++ and resuspended in 10% FBS in PBS for 30 min to block non-specific staining. Cells were then pelleted by centrifugation and incubated with primary antibody on ice for 30 min. Following three rounds of washing with PBS by centrifugation, cells were incubated with AF647-conjugated secondary antibodies for 30 min on ice, washed twice and filtered through 50μm filters. Samples were analysed on a Fortessa flow cytometer and further analysis undertaken using FlowJo software.

Flow cytometric sorting of single-cell dissociated organoids transduced with fluorescent markers was performed using a sorter (BD FACSMelody) and analysed using FlowJo.

### Immunoblotting

For immunoblotting, cells were subjected to triton lysis, SDS-PAGE electrophoresis, and immunoblotting as previously described^17^. Membranes were incubated overnight with primary antibody at 4°C before washing and incubating with secondary antibody (1:20,000 IRDye® conjugated, various) for 1 hour at room temperature. Membranes were visualised using a Li-Cor Odyssey imaging system.

### Immunofluorescence staining and microscopy

For paraffin embedding, organoids were seeded onto 0.4μm transwell inserts embedded in 50% matrigel. Organoids were fixed with 4% PFA for 30 min before the membrane was released from the insert using a scalpel and embedded between layers of HistoGel before paraffin embedding. Samples were deparaffinised and rehydrated using sequential passes through xylene (x3) then ethanol (100%, 70%, 50%, 0%) then antigen retrieval undertaken by boiling slides in sodium citrate buffer (10mM sodium citrate, 0.05% tween 20 pH 6.0) for 20 min. Samples were blocked using 1% BSA, 0.1% tween then incubated with primary antibodies overnight and Alexa-fluor conjugated secondary antibodies (1:500, various, ThermoFisher) for 1 hour at room temperature before staining with DAPI, and mounting with ProLong Gold Anti-fade Mountant.

Wholemount staining was performed on fdAT2 organoids grown in Nunc Lab-Tek II chamber slides (Thermo, 154453) and fdAT2 monolayers in transwell inserts. Samples were washed in PBS before fixation with 4% PFA in PBS for 30 minutes at room temperature. Samples were then washed and incubated with NH_4_Cl (50mM in PBS) for 30 minutes at room temperature to quench autofluorescence before permeabilisation with 0.2% Triton X-100 in PBS for 30 minutes at room temperature. After blocking with 10% FBS in PBS for 60 minutes, samples were incubated with primary antibody overnight at 4°C, washed, and incubated with secondary antibody for 2 hours at room temperature.

Samples then underwent 3 x 5 minute wash with PBS, with DAPI added at a concentration of 1µg/mL for the penultimate wash. Chambers were removed from slides before mounting coverslips with ProLong Gold Antifade. Images were taken using a Zeiss LSM880 with airyscan and Zen black software.

For lysosomal fluorescence, fdAT2 grown either as a monolayer on embedded in matrigel were incubated with 50nM Lysotracker in fdAT2 medium at 37°C for 1.5 hours. For embedded cultures, organoids were removed from the matrigel with cold media, isolated by centrifugation and placed directly into an ibidi imaging chamber slide. For monolayer cultures, membranes were excised and transferred to a live cell imaging dish for immediate imaging.

For cell viability assays, fdAT2 grown as a monolayer were incubated with Hoescht 33342 diluted 1:10,000 and 7-AAD diluted 1:40 in fdAT2 medium for 10 minutes then washed with warm DMEM+++. Membranes were excised and transferred to a live cell imaging dish for immediate imaging.

### Wound healing assays

6-well transwell inserts were coated with 20% matrigel before 2x 3-well culture-inserts were attached to each membrane. Dissociated fdAT2 were plated into the 3-well inserts at a density of ∼2×10^5^ cells per well, with WT/WT and WT/I73T cells placed in separate inserts on the same transwell. Once confluent (48hrs), cells were air-exposed or kept submerged before wounding by removal of the insert 24hr later. Four images per condition were taken on alternate days using an axioobserver, with intra-image reference points used to ensure the same field of view was captured. Wound healing was calculated by delineating the remaining wound area in each image using Zen blue and calculating the mean for each experiment (n=3 independent experiments on one biological line). Light intensity measurements were plotted using Zen blue and the mean of 4 cross sections reported.

### Quantitative RT-PCR

Total RNA was isolated using an RNeasy kit including an optional DNase digestion step. Typically 500ng RNA was used as the starting template to create cDNA using a high capacity cDNA reverse transcription kit and heating samples to 25°C for 10 min, 37°C for 2 hours and 85°C for 5 min. RT-qPCR was undertaken in 96 well plates using 4.5μl 1:10 cDNA and 10.5μl of a master mix containing SYBR Green JumpStart Taq Readymix. Primer sequences are listed in table 3. Plates were run on a BioRad RT PCR machine typically using the following programme: 95°C 2 min, 40x (95°C for 30 sec, 55°C for 30 sec, 72°C for 30 sec), 95°C for 30 sec.

**Table 3:**
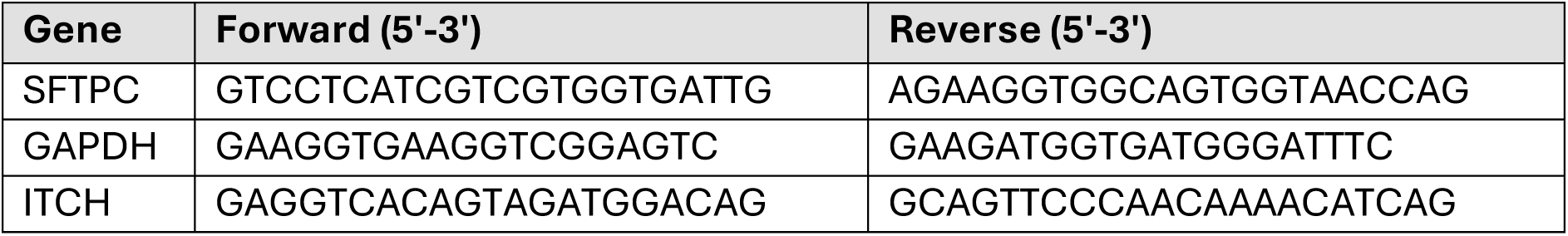
qPCR primers.

### Electron microscopy

Whole organoids were fixed with 2% PFA, 2.5% glutaraldehyde, 0.1M cacodylate buffer, pH 7.4. Organoids were secondarily fixed with 1% osmium tetroxide/1.5% potassium ferrocyanide and then incubated with 1% tannic acid in 0.1M cacodylate buffer to enhance membrane contrast. Organoids were washed with water before being dehydrated using increasing percentages of ethanol (70,%, 90%, 100%). Samples were embedded in beam capsules in CY212 Epoxy resin and resin cured overnight at 65°C. Ultrathin sections were cut using a diamond knife mounted to a Reichart ultracut S ultramicrotome. Sections were collected onto piloform-coated slot grids and stained using lead citrate. Sections were viewed on a FEI Tecnai transmission electron microscope at a working voltage of 80kV.

### Statistical analysis

Statistical analysis and data visualisation were performed using Graphpad Prism software (version 9.5.1). Data from at least three independent experiments using three biologically independent lines (unless specified otherwise) was used for statistical analysis and is expressed as mean ± standard deviation (SD). Details of statistical tests used are detailed in figure legends. Lysotracker-positive vesicles were measured and cell viability quantified (% of dual Hoecht and 7-AAD positive cells) using Imaris software. Measurement of EEA1 positive vesicles was undertaken manually as automated methods were unable to adequately delineate structures.

---

**Suppl Fig 1.**
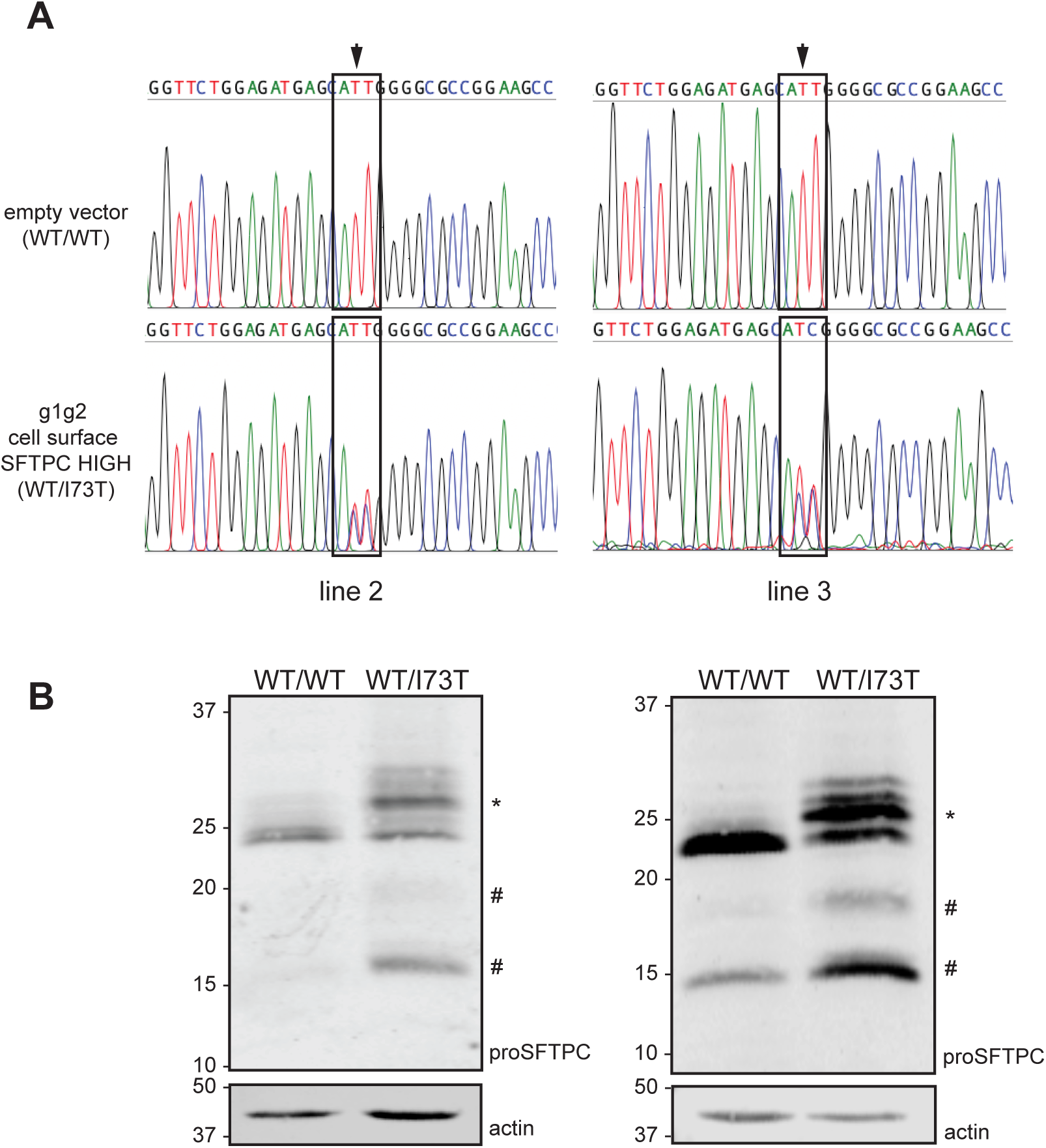
Characterisation of n=3 base edited fdAT2 lines. (A) Sanger sequencing of lines 2 and 3 following transduction with base editing guides and sorting for cell surface SFTPC enriched cells. Relevant codon (ATT-ACT) highlighted. (B) proSFTPC immunoblotting of SFTPC-WT/WT and WT/I73T fdAT2. *higher molecular weight full length SFTPC-I73T caused by O-glycosylation; #partially and fully C-terminally cleaved intermediates.

**Suppl Fig 2.**
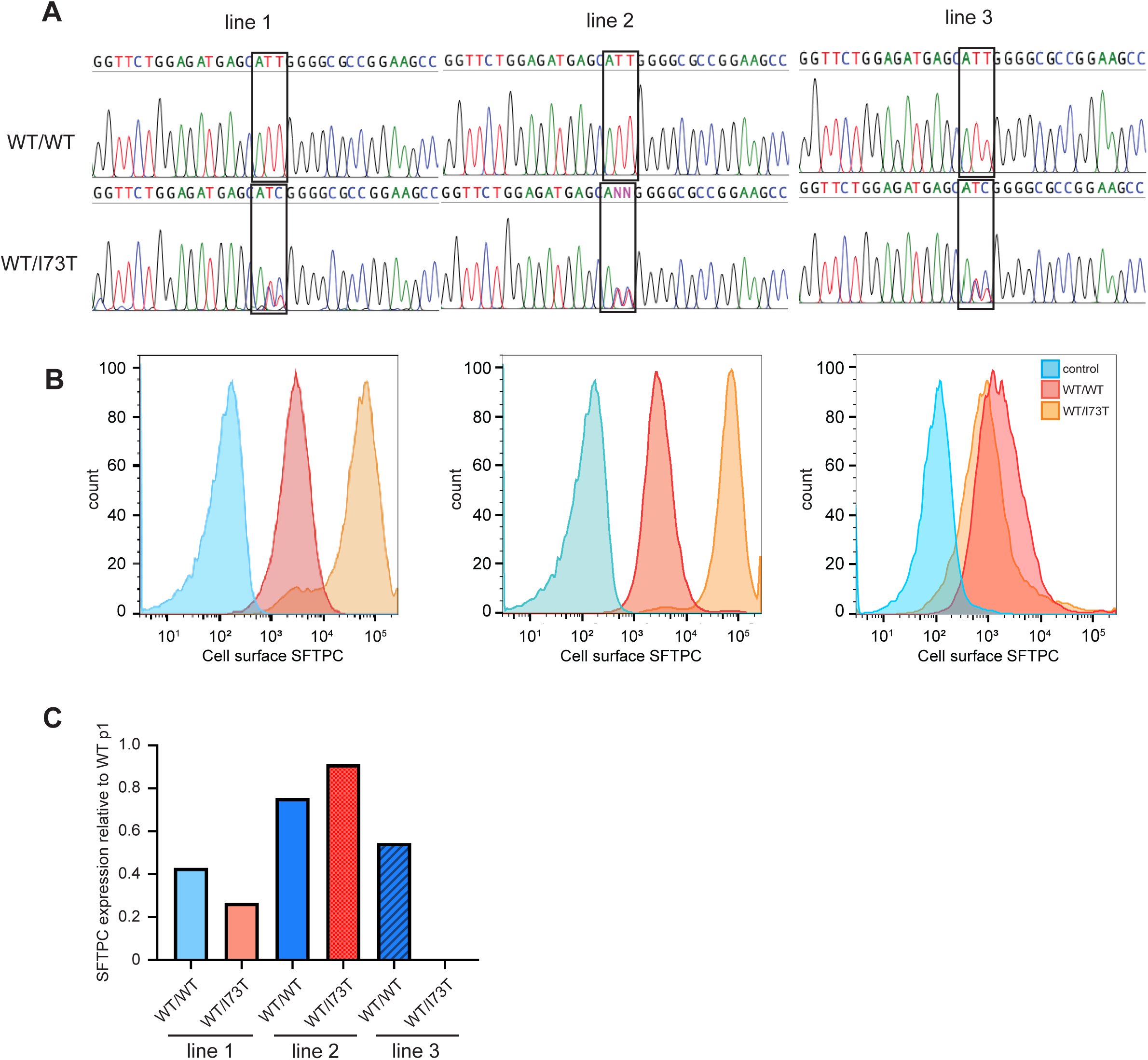
Genotypic and phenotypic characterisation of later passage edited fdAT2 lines. (A) Sanger sequencing of n=3 biologically independent fdAT2 lines at passage 10. (B) Flow cytometry histogram of cell surface SFTPC in the 3 lines as measured by C-terminal domain antibody. (C) SFTPC expression in passage 10 fdAT2 lines. Expression relative to GAPDH and normalised to expression in equivalent WT/WT line at p1 (n=1 in triplicate).

**Suppl Fig 3.**
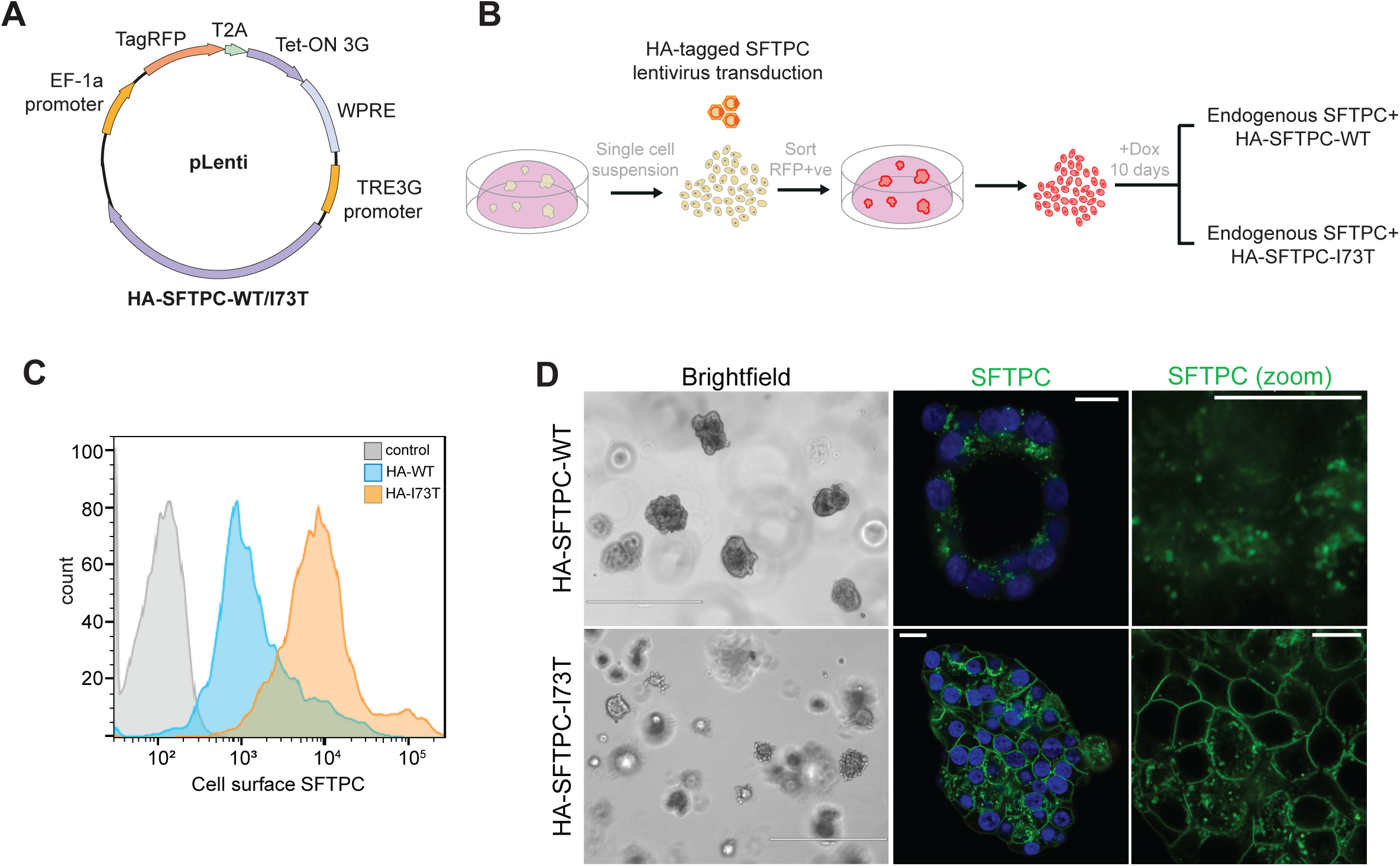
Generation of HA-SFTPC fdAT2 lines. (A) pLenti expression plasmid containing HA-SFTPC and EF-1a-RFP to allow fluorescence sorting. (B) Schematic of HA-SFTPC fdAT2 generation. (C) Example flow cytometry histogram of cell surface SFTPC as measured by C-terminal domain antibody after 10 days of induction. (D) Brightfield imaging and immuofluorescence imaging of SFTPC localisation in fdAT2 expressing HA-SFTPC-WT or I73T for 10 days. Scale bars, 400µm (brightfield), 10µm (immunofluorescence).

**Suppl Fig 4.**
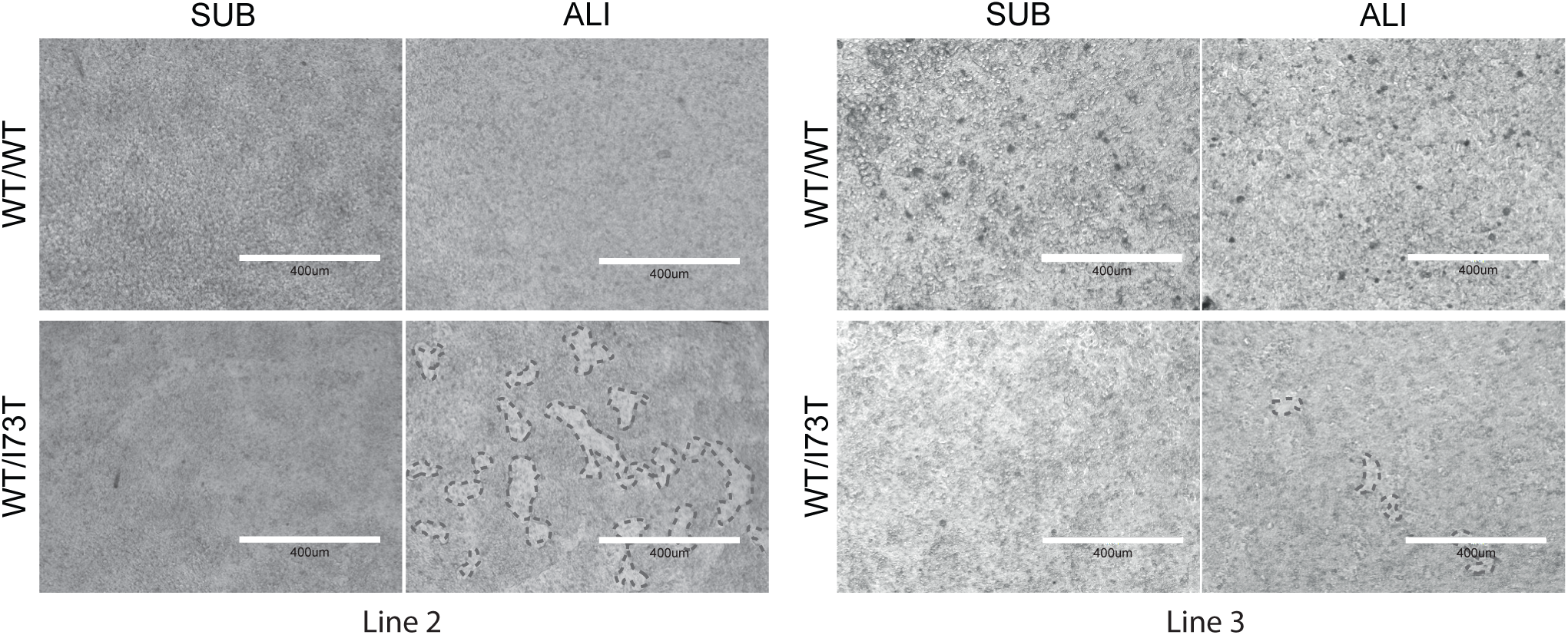
Transwell culture phenotypes of n=3 edited fdAT2 lines. Brightfield imaging of fdAT2 monolayer in lines 2 and 3; discontinuities demarcated with dotted lines. Scale bar, 400µm.

**Suppl Fig 5.**
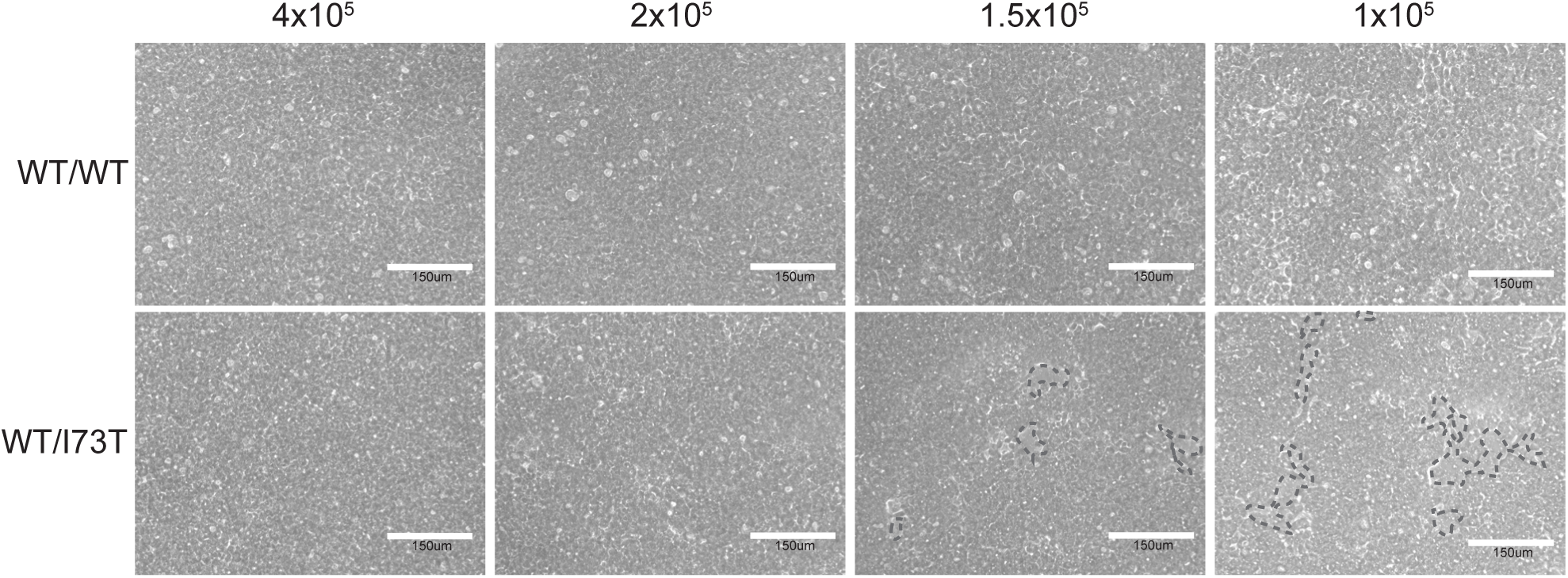
Seeding density optimisation for wound healing assays. fdAT2 seeded at varying densities and air-exposed for 48 hours to assess monolayer integrity. Cell number = total for 24-well insert. Scale bar, 150µm.

